# Reversing Radiation-Induced Immunosuppression Using a New Therapeutic Modality

**DOI:** 10.1101/2022.05.03.490472

**Authors:** Colleen M. Courtney, Sadhana Sharma, Christina Fallgren, Michael M. Weil, Anushree Chatterjee, Prashant Nagpal

**Affiliations:** Sachi Bioworks, 685 S Arthur Avenue, Colorado Technology Center, Louisville, CO 80027; Environmental & Radiological Health Sciences, Colorado State University, Fort Collins, CO 80523

**Author notes:** Corresponding Author., 685 S. Arthur Ave, Unit 5, Louisville, CO 80027.

**Keywords:** Radiation-induced immunosuppression, immune modulation, gene therapy, Nanoligomer, drug-discovery, target validation

## Abstract

Radiation-induced immune suppression poses significant health challenges for millions of patients undergoing cancer chemotherapy and radiotherapy treatment, and astronauts and space tourists travelling to outer space. While a limited number of recombinant protein therapies, such a Sargramostim, are approved for accelerating hematologic recovery, the pronounced role of granulocyte-macrophage colony-stimulating factor (GM-CSF or CSF2) as a proinflammatory cytokine poses additional challenges in creating immune dysfunction towards pathogenic autoimmune diseases. Here we present an approach to high-throughput drug-discovery, target validation, and lead molecule identification using nucleic acid-based molecules. These Nanoligomer™ molecules are rationally designed using a bioinformatics and an artificial intelligence (AI)-based ranking method and synthesized as a single-modality combining 6-different design elements to up- or downregulate gene expression of target gene, resulting in elevated or diminished protein expression of intended target. This method additionally alters related gene network targets ultimately resulting in pathway modulation. This approach was used to perturb and identify the most effective upstream regulators and canonical pathways for therapeutic intervention to reverse radiation-induced immunosuppression. The lead Nanoligomer*™* identified in a screen of human donor derived peripheral blood mononuclear cells (PBMCs) upregulated Erythropoietin (EPO) and showed the greatest reversal of radiation induced cytokine changes. It was further tested *in vivo* in a mouse radiation-model with low-dose (3 mg/kg) intraperitoneal administration and was shown to regulate gene expression of *epo* in lung tissue as well as counter immune suppression. These results point to the broader applicability of our approach towards drug-discovery, and potential for further investigation of lead molecule as reversible gene therapy to treat adverse health outcomes induced by radiation exposure.

## INTRODUCTION

In space exploration missions or travel to outer deep space, astronauts and space tourists are subjected to increased amounts of radiation exposure, which can lead to Acute Radiation Syndrome (ARS) causing DNA damage, tissue damage, and aging.^1^ ARS or radiation toxicity is an acute illness caused by radiation exposure on a part of or whole body by a high dose (>1 Gray or Gy) of radiation for a short period of time.^2, 3^ Classic ARS syndromes include hematopoietic-ARS (H-ARS) and Gastrointestinal-ARS (GI-ARS), which typically occur with cumulative radiation dose between 0.7 and 10 Gy. As space missions get longer and farther away from the lower Earth orbit and thus from Earth’s protecting magnetic shielding field, the radiation exposures that astronauts or space tourists face expand to the full galactic cosmic ray spectrum and solar particle events. Ionizing radiation (including gamma or γ-rays) is a ubiquitous and dangerous environmental hazard during deep and long-term space explorations and is associated with immune dysfunction that can lead to development of cancers and increased susceptibility to infectious diseases and latent viruses.^4–8^ These health threats are also faced by unsuspecting civilian populations in case of accidental or malicious nuclear incidents.^9^ Similar effects are faced by patients undergoing cancer chemotherapy and radiotherapy, where the radiation-induced immune suppression makes the patients immunocompromised, susceptible to infections, and leads to poor clinical outcomes.^10, 11^

The radiation countermeasures currently available for treatment of ARS and H-ARS are very limited.^3^ H-ARS is driven by loss of crucial growth factor-modulated hematopoietic progenitors and consequently, by large losses of circulating, functional blood cells. This condition has been shown to be treated by recombinant colony stimulating factors (CSFs)^5, 11–14^, specifically granulocyte-colony stimulating factor (G-CSF or CSF3)^5, 11, 13, 14^ and granulocyte-macrophage colony-stimulating factor (GM-CSF or CSF2).^5, 11–14^ CSF3 is a glycoprotein produced by monocytes, fibroblasts, and endothelial cells that induces bone marrow hematopoietic progenitors to differentiate into specific mature blood cell types released into the bloodstream. CSF2 is a monomeric glycoprotein that functions as a cytokine and is secreted by macrophages, T-cells, mast cells, natural killer cells, endothelial cells and fibroblasts. To date, there have been several studies involving radiation accidents where ARS patients were treated with recombinant CSF3 and CSF2 with limited success.^15–17^ Other proteins playing a role in radiation response include Erythropoietin (EPO), a glycoprotein cytokine which is secreted by the kidney in response to cellular hypoxia. EPO stimulates red blood cell production (erythropoiesis) in the bone marrow and treatment with recombinant EPO has been shown to induce cancer cell resistance to ionizing radiation and cisplatin, and thus is considered a possible radiation countermeasure.^18, 19^ One of the biggest challenges is that these therapeutics are provided as recombinant proteins (such as FDA approved Leukine (Sargramostim), Neupogen (Filgrastim), and Neulasta (Pegfilgrastim)), and while they alleviate some of the phenotypic biomarkers caused due to ARS and H-ARS, several multi-omics and radiobiological studies have indicated need for inclusion of other targets to provide more effective radiation-exposure treatment.^5, 11, 12–14, 18–22, 23^ These therapies are approved as radiation treatment countermeasures to lessen the effects after radiation exposure and onset of ARS symptoms. Some other practical challenges for space applications involve long-term production and storage on spaceflight which is complicated for recombinant proteins. Recent research efforts using nucleic acid therapy of mammalian cells include microRNAs (miRNA), small interfering RNAs (siRNA), long non-coding RNAs (lncRNA), and CRISPR technology; though promising, these therapeutic strategies involve tedious cloning and optimization that are time-consuming. At present, nucleic acid-based therapies do not exist for any of the above targets. Furthermore, recent studies have shown that there are other key players involved in radiation associated damage and targeting them using conventional approaches such as recombinant proteins or small molecule drugs is time consuming, laborious, expensive and ultimately not feasible. ^2, 3, 5, 11–14, 18–20, 21–30^ To address the limitations of the existing approaches, we have developed a new approach to high-throughput drug-discovery, target validation, and lead molecule identification using nucleic acid-based molecules, called Nanoligomer™, that are rationally designed and synthesized to up- or downregulate expression of target gene at the messenger RNA (mRNA) or protein level. Furthermore, it alters related gene network targets ultimately resulting in pathway modulation.

Recent studies have performed high throughput gene expression analysis in tumor cells and cell lines, primary cells, rodents, and peripheral blood of patients to accurately identify key genes involved in radiation response.^28, 31^ These studies highlighted the importance of multi-gene response to radiation, specifically those genes involved in natural kill cell activation, B-cell mediated immunity,^27^ and gene regulated by three known radiation-modulated transcription factors tumor protein 53(TP53), Forkhead Box M1 (FOXM1), and erb-b2 receptor tyrosine kinase 2 (ERBB2).^28^ Additionally, the most comprehensive set of omics-data was recently published through the NASA twin study, providing unprecedented insights into the regulatory cytokine and proteins involved in repair and recovery by the body as countermeasures following a year-long space mission. This study showed the importance of cytokines such as Tumor Necrosis Factor alpha (TNF-α), type I interferons (IFN), GM-CSF, Interleukin 4 (IL-4), Interleukin 5 (IL-5), Interleukin 6 (IL-6), Interleukin 10 (IL-10), Interleukin 12 (IL-12), Interleukin 18 (IL-18), vascular endothelial growth factor (VEGF), basic fibroblast growth factor (bFGF) and transforming growth factor beta (TGF-β) as markers of radiation exposure recovery.^5^ In monotherapeutic and combination therapy of stem-cell factor (SCF), FMS-like tyrosine kinase 3 ligand (FLT-3 ligand), thrombopoietin (TPO), Interleukin-3 (IL-3) and stromal derived factor-1 (SCF-1) in mice under acute radiation exposure (8 Gy), it was shown that the survivability increases from 8.3% under no treatment to ∼29% for individual therapy, to 87.5% for combination therapy.^12^ While these statistics reveal the potential for these combination proteins in ensuring short-term survival (30 days), their long-term survival was lower, presumably dependent on the half-life of injected proteins and combinations. These studies and other radiobiological data, ^3, 5, 11–14, 18–21, 22, 27, 28, 30^ provide reproducible and reliable immunological targets identified across independent studies to treat pathology induced due to radiation exposure, included as targets in our study (Table 1). Here, we apply our Nanoligomer platform to perturb selected immunological targets (Table 1) to identify most effective upstream regulators and canonical pathways for therapeutic intervention to reverse radiation-induced immunosuppression.

**Table 1:**
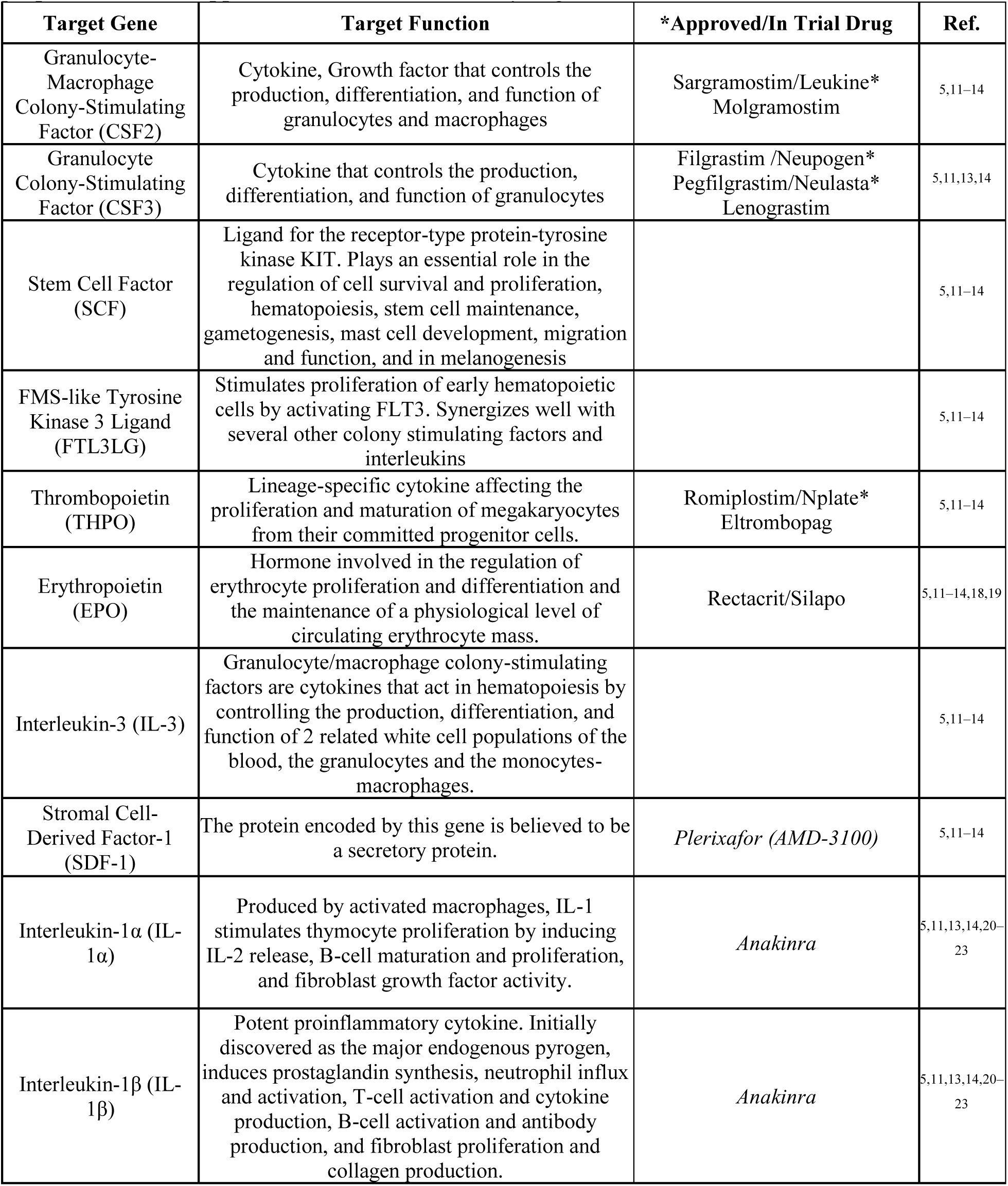
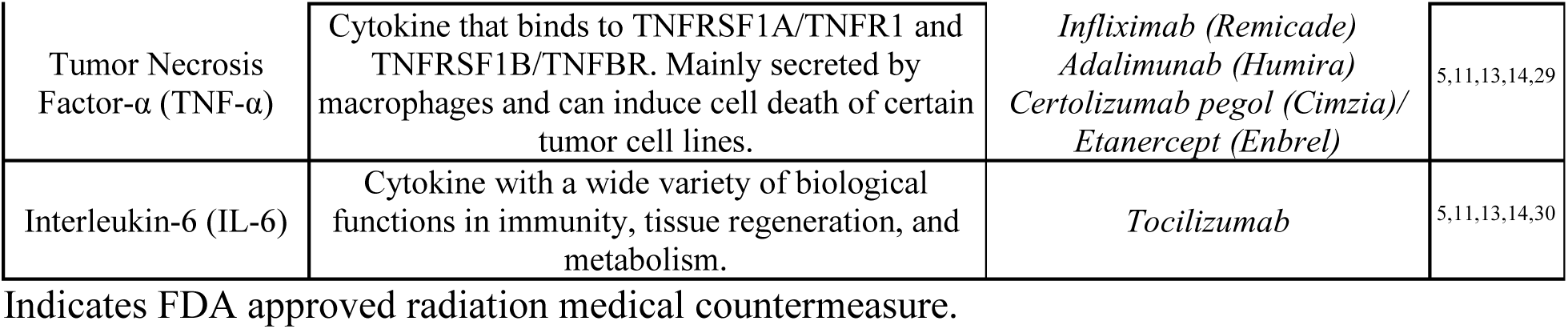
Prioritized gene targets for radiation medical countermeasure development. Italics of Approved/In Trial Drug indicates that the mechanism of action of the drug on target gene is proposed to be the opposite needed for immune dysregulation correction

## Results and Discussion

The Sachi platform presented here generates sequence-specific, nano-biohybrid therapeutic candidates called Nanoligomers^TM^ (Sachi Bioworks) (Fig.1). The nucleic acid-binding domain of Nanoligomers is peptide nucleic acid (PNA), which is a synthetic DNA-analog where the phosphodiester bond is replaced with 2-N-aminoethylglycine units. PNA demonstrates strong hybridization and target specificity compared to binding of naturally occurring RNA or DNA,^32, 33^ as well as exhibits no known enzymatic degradation,^32, 33^ leading to increased stability in human blood serum and mammalian cellular extracts.^34, 35^ However, PNAs as well as other antisense therapeutics suffer from transport challenges.^36^

**Fig.1.**
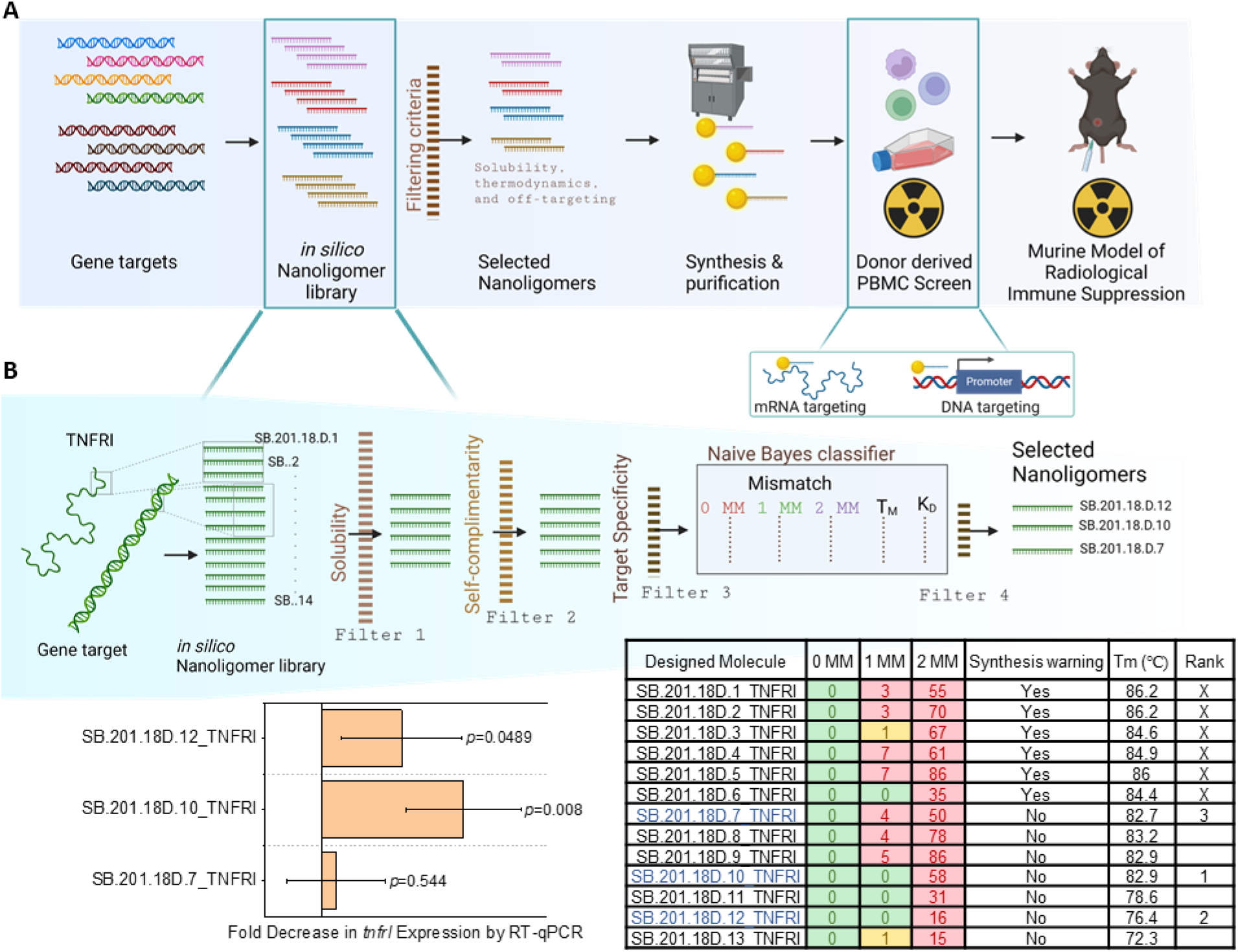
Evaluation of Nanoligomers on radiological immune suppression in human donor derived PBMCs and mouse. **A)** For radiological immune regulation, an *in silico* Nanoligomer library was created from chosen gene targets. Using our pipeline, we constructed a list of prioritized candidate Nanoligomers which were synthesized and purified in our platform with functionalized nanomaterials. These Nanoligomers will first be screened in irradiated donor derived PBMCs to evaluate the effect on radiological immune response via 65-plex cytokine panel. Finally, an identified top Nanoligomer candidate will be tested in a murine model of radiological immune suppression in lung and evaluated by cytokine panel for its ability to reduce radiological immune suppression. **B)** Our *in silico* Nanoligomer library was created (shown here with sample gene *tnfr1* (tumor necrosis factor receptor 1) and candidates evaluated and selected through our four-step process. First, gene target regions (RNA and/or DNA) are identified based on factors such as sequence composition and predicted transcription or translational regulatory regions. These seed regions are then evaluated for expected synthesis and solubility considerations followed by filtering based on undesirable features such as self-complementary and direct match or multiple mismatch off-targets. The final step is a thermodynamic analysis using a Naïve Bayes classifier for target vs off-target binding and optimization of possible on-target effectiveness. Three Nanoligomers were tested from sample gene *tnfrI* and varied effectiveness was measured on gene expression which correlated with our ranked *in silico* selection of top Nanoligomer.

Sachi’s proprietary Nanoligomer molecule employs an engineered nanoparticle to remove bottlenecks in product synthesis/purification and achieve improved cellular and tissue delivery. These features make Nanoligomers especially attractive candidates for both interrogating targets and developing nucleic acid therapies. Nanoligomers are designed to bind to RNA (mRNA, miRNA, lncRNA, etc.) or DNA (Fig.1). Gene expression activation is achieved by building activators that target binding at the respective promoter regions and have specific domains attached that recruit transcriptional activators, thereby causing transcriptional activation. RNA inhibition is achieved by blocking translation of targeted mRNAs or blocking functional regions of non-coding RNA. Additionally, RNA inhibition can be achieved by signaling for RNase degradation of the target RNA.

The platform utilizes a bioinformatics toolbox to design a Nanoligomer that will selectively downregulate or upregulate the gene of interest while minimizing off-targeting and optimizing thermodynamic properties of the molecule (Fig.1B). The bioinformatics toolbox performs off-targeting analysis for up to two base pair mismatches to the human genome thereby promoting specificity of Nanoligomers to their intended target gene. Further, the software optimizes for binding affinity, solubility, and other thermodynamic parameters in effort to maximize therapeutic efficacy.

### Scoring and Ranking Bioinformatic Generated in silico Nanoligomer Library

We utilize machine-learning based approaches and experimental data to rank, and then validate, the different candidates for gene-perturbation of the potential target (see Methods, Table S1). During initial platform development, we utilized experimentally measured/calculated biochemical data (for example *K_D_* (disassociation constant), *T_M_* (melting temperature of binding event)) to set up the scoring method, and then utilized the experimental gene-perturbation measurements from different candidate Nanoligomers (Fig.1B) to set up: a) training set; and b) testing set, and then iterate between the training and testing sets to optimize the model development.^37–42^ To validate and rank the candidates in this study, we utilized the model to design the lead Nanoligomer for each target and then used it to the assess the relative importance of multiple targets (lead target ID) in Table 1.

### Assessing Relative Importance/Efficacy of Target in Reversing Immunosuppression

The lead Nanoligomer molecule for each target was identified using the platform (Fig.2A), synthesized, and then screened using human donor-derived peripheral blood mononuclear cells (PBMCs), a well-used *in vitro* test platform for immune modulation,^43, 44^ with and without radiation exposure. In addition to the Nanoligomer library, we tested three small-molecules that have been studied as radiological countermeasures as comparison in our assays: 100 µM Aspirin (Asp, medication used to reduce pain, fever, or inflammation and has demonstrated reduced radiation toxicity)^45^; 10 µM Metformin (MetF, anti-diabetic medication; controls high blood sugar),^46^ and 100 µM Epicatechin (EC, ROS scavenger as radioprotective countermeasure).^47^ PBMC were treated with either chemical inhibitor or 10 µM Nanoligomer and incubated for 24 hours. Following incubation, cells were irradiated, allowed to recover for 4 hours, and centrifuged to collect supernatant for cytokine analysis and cell pellet for RNA analysis.

**Fig.2.**
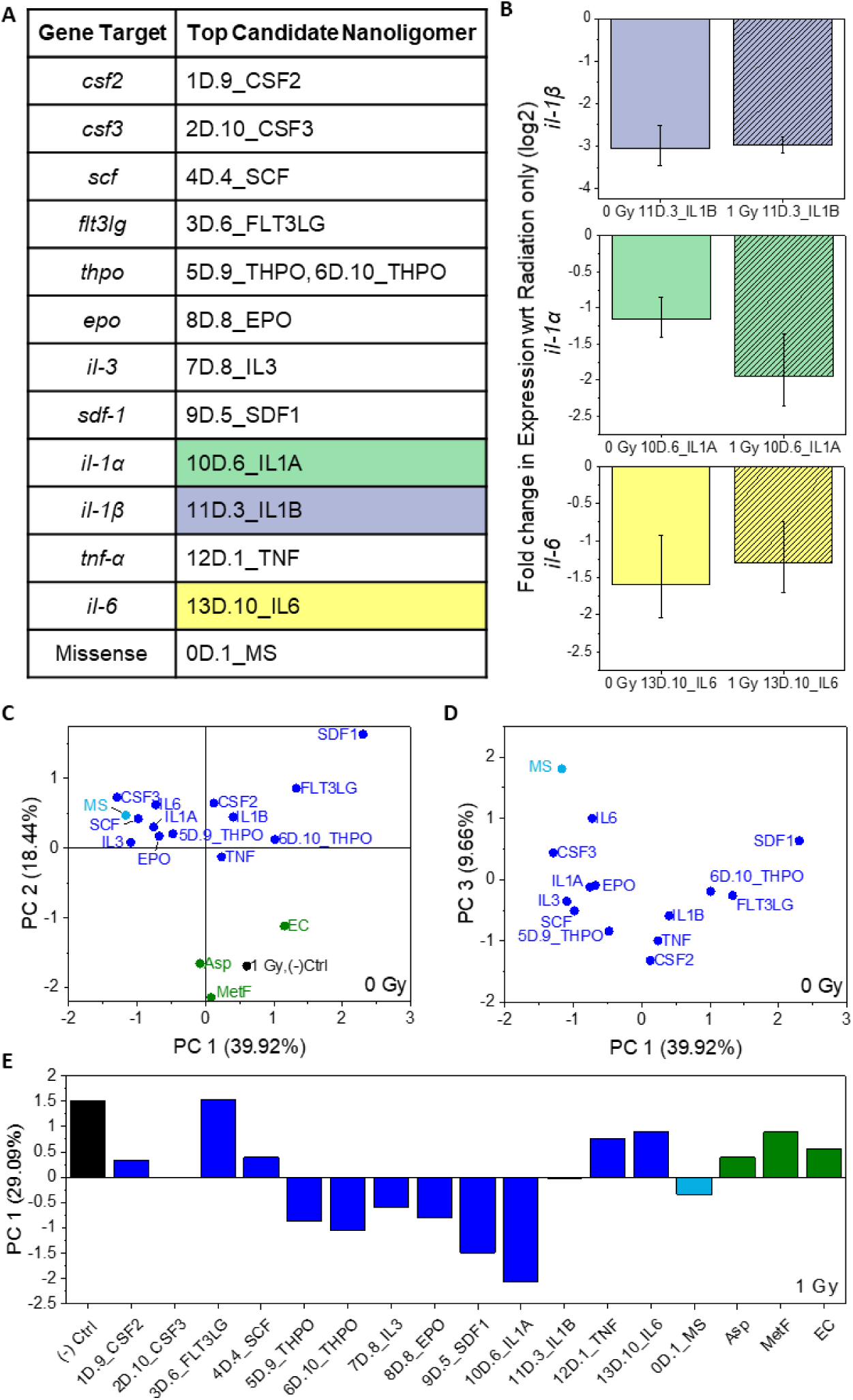
Candidate Nanoligomers and Gene Expression for three selected treatments and corresponding target. **A)** Library of candidates where “D” indicates a downregulator. **B)** Gene expression validation wrt no Nanoligomer treatment (negative control) for PBMCs. Each is normalized to it’s no Nanoligomer with respective irradiation. (Top) more than 8- fold reduction in IL-1B expression, at 0 and 1 Gy, compared to their untreated PBMCs irradiated with same dosage. (Middle) More than 2-3-fold decrease in IL-1A expression due to Nanoligomer treatment. (Bottom) More than 3-fold change in IL-6 gene expression observed for human PBMCs. **C-D)** Human donor derived PBMC were treated with Nanoligomers (here data labeled by their gene target), incubated for 24 hours and collected for cytokine 65-plex panel analysis. Principal component statistical analysis was performed on the result to elucidate distinctions between treatment groups. Included in this analysis was also No treatment with 1 Gy radiation for comparison of an irradiated cytokine signature. PC1 vs. PC2 (left) demonstrates that the chemical treatments (Epicatechin (EC), Aspirin (Asp), and Metformin (MetF)) look similar in profile to an irradiated PBMC sample while the Nanoligomers are difference from 1 Gy radiation with no treatment. PC1 vs PC3 shows signatures that separates the missense Nanoligomer from the rest of the gene targeting Nanoligomers. **E)** Human donor derived PBMC were treated with Nanoligomers, incubated for 24 hours, irradiated with 1 Gy, and collected for cytokine 65-plex panel analysis. Principal component analysis highlights the statistical difference between many Nanoligomer treatments and the no treatment negative control.

In a representative set, we validated that the Nanoligomers indeed perturb the gene expression of their intended target using 10D.6_IL1A, 11D.3_IL1B, and 13D.10_IL6 Nanoligomers (Fig.2B, S1A). Off-target effects were investigated using multiplexed analysis of non-pathway genes and indicated gene target specificity (Fig.S1B). RNA samples were prepared from PBMCs for both non-irradiated 0 Gy and radiation-treated 1 Gy samples and RNA quantification was performed using Thermo Fisher Scientific QuantiGene (see methods). All three Nanoligomers showed significant downregulation of the target with and without radiation.

The collected supernatant was analyzed using high-throughput multiplexed Thermo Fisher Scientific ProcartaPlex for a comprehensive panel of 65-human cytokines, chemokines, growth factors, and other biomarkers of immune response (see methods). Data was normalized with respect to non-treated PBMCs (No trt, Fig.S2) and the high-dimensional (65-plex) data was analyzed using principal component analysis (PCA). PCA enables complex data sets to be simplified into fewer dimensions (i.e., components) to find trends and patterns.^48^ Additionally, PCA is non-bias, like clustering, and can summarize effects within the data set with less noise allowing pattern recognition and data summary. Each principal component (PC) is calculated to be geometrically orthogonal and uncorrelated with all other PCs to provide simplified, distinct dimensions. Most usable trends and patterns were comprised in the first few PCs and their strength as a component was determined by the percent of variability contained in the component and by examining Scree plots of all calculated PCs (Fig.S5). Pitfalls of PCA stem from the need to have data on a similar scale (i.e., higher magnitude values will dominate PC compared to smaller magnitude values) and its treatment of the data as linear and uncorrelated. For analysis, all data in PCA was normalized to the corresponding cytokine level in the negative control to minimize issues of data scaling.

The cytokine, chemokine, growth factor/regulators, and other soluble receptor data from 65-biomarkers in human immune monitoring panel was assessed with the 14 Nanoligomer treatments (Fig.S3), three chemical inhibitors (Fig.S4), and analyzed with untreated PBMCs exposed to 1 Gy gamma radiation (Fig.S2). The PCA Scree plot (Fig.S5) shows the importance of the first three principal components (PC1, PC2, and PC3). These PCs account for 39.92%, 18.44%, and 9.66% of variance, respectively, and additionally include key cytokines identified as top/important biomarkers in the immune dysregulation in the NASA twin study (Fig.2C, S6, Table S2). PC3 provides further distinction between all Nanoligomers tested including high separation of the missense from targeted Nanoligomers (Fig.2D). Using the scores assigned with these PCs (score plot, Fig.2C, 2D), there were several important quantitative (as numerical scores) and qualitative assessment parameters obtained. Briefly, 1) 4 key targets were identified as being the lead gene targets in reversing immune dysregulation as those farthest from the untreated (No Trt) samples at 1 Gy (Fig.3A, EPO, CSF2, CSF3, IL3); 2) Some other Nanoligomers such as 6D.10_THPO, 11D.3_IL1B, etc. showed variance in many key cytokines and were further away from 1 Gy, no treatment, but scored less strongly as 4 lead targets; and 3) chemical inhibitors showed similar characteristics as 1 Gy, no treatment for immune dysfunction, and likely had other orthogonal mechanism of radioprotection not captured using the *in vitro* immune model used here. The lead targets were then evaluated using the same PBMC model but now irradiated at 1 Gy (Fig.S7) and 2 Gy (Fig.S8) gamma-irradiation. Using only the first PC which contained 29.09% of variability to quantitatively assess and rank the selected targets (as evaluated using their Nanoligomers), we identified the following ranking for top five targets at 1 Gy: SDF, THPO-iso, EPO, IL3, and tied CSF2 and CSF3 (Fig.2E). Once we determined a small set of key Nanoligomers, we analyzed the panel data for specific changes in cytokine expression. First, any cytokine that showed broad expression changes across multiple treatments were removed from analysis to elucidate gene specific effects and highlight unique and top Nanoligomers. Cytokine levels were normalized to no treatment with 2 Gy radiation and significance was calculated using a t-test between replicates (Fig.3). This evaluation at 2 Gy revealed the top target as *epo* based on significant changes by 8D.8_EPO Nanoligomer in relevant cytokines, growth factors/regulators, and chemokines for immune dysregulation.

**Fig.3.**
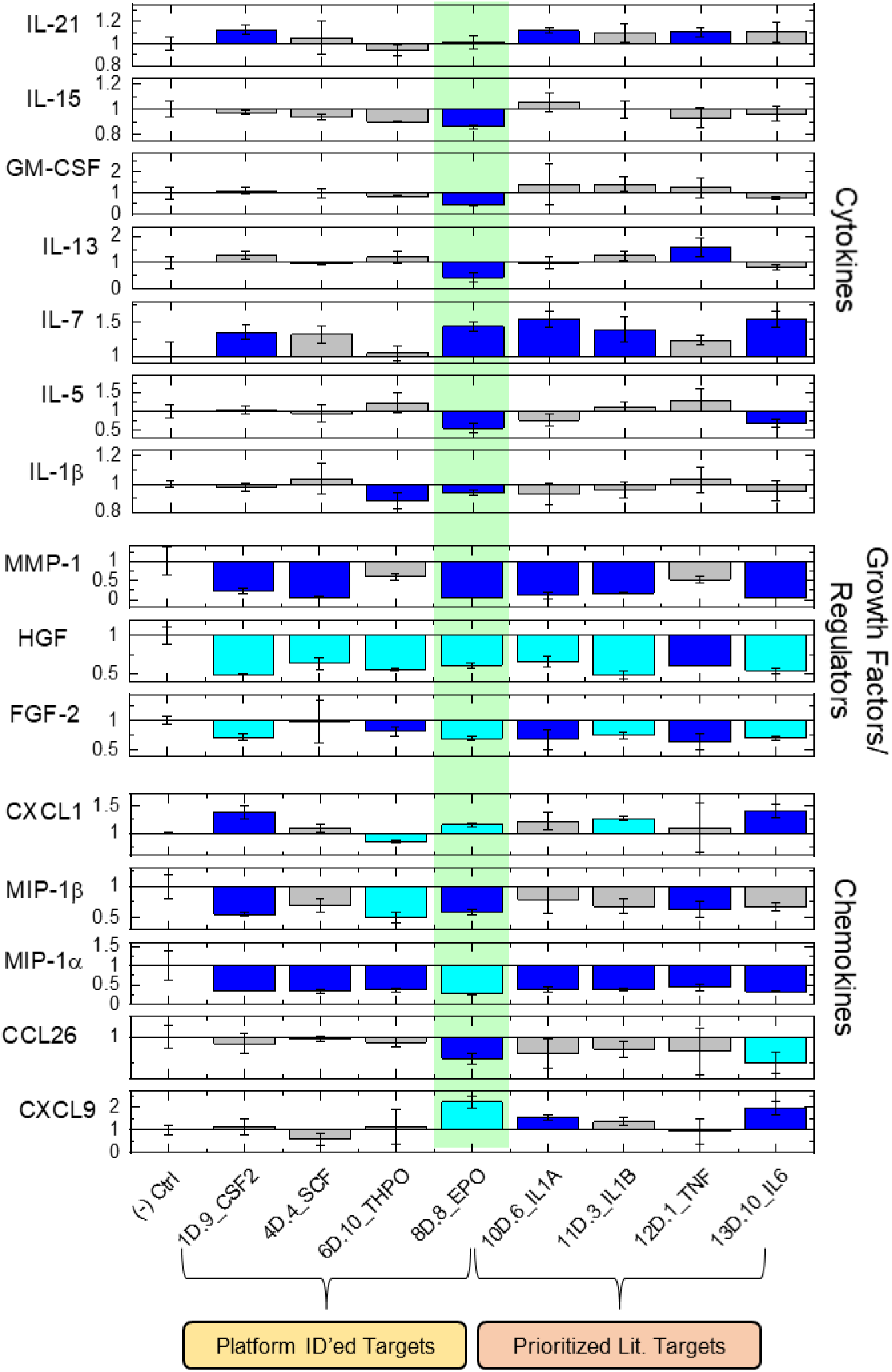
Nanoligomers induced significant changes in cytokine, growth factors/regulators, and chemokines with 2 Gy radiation in human PBMC under radiation induced immune dysregulation. The down selected Nanoligomers were again screened in higher total radiation dose of 2 Gy in human donor derived PBMC. Fold change of protein in PBMC supernatant demonstrates the effect of Nanoligomer where less than 1 is less protein in treated, equal to 1 is similar to untreated, and greater than 1 is more protein in treated cells. Any cytokine that showed a change with missense Nanoligomer was not included for this analysis to rule our Nanoligomer specific changes. Blue color represents *p*<0.05 and cyan color represents *p*<0.01 with respect to (-) Ctrl (2 Gy, PBS). The green shading shows our top target gene identified by Nanoligomer downregulator screen, EPO.

### Nanoligomer *epo* Upregulator Demonstrates Reversal of Immune Suppression and Increased Regulatory T Cell Populations

Using our pipeline, we next designed an upregulator for *epo* (8U.1_EPO) as well as an upregulator for *csf2* (1U.2_CSF2) because of its demonstrated use as a radiological countermeasure. We additionally synthesized a missense control upregulator (0U.1_MS). These Nanoligomers were then tested in PBMCs with total radiation dose of 3 Gy and the supernatant was analyzed by 65-plex ProcartaPlex panel (Fig.S9). PCA performed on the data and examination of principal component 1 and 2 (PC1 and PC2) which account for 73.62% and 11.31% of variability, respectively, highlights a distinction between no treatment, missense treated cells, and targeted Nanoligomers to *epo* and *csf2* (Fig.4A). The 8U.1_EPO and 1U2._CSF2 treated cells had similar cytokine profiles, which we hypothesize is likely due to the closely related gene network of the two targets. Notably, both upregulators caused significant changes in cytokines known to be important for radiation induced immune dysfunction (Fig.4B). Nine of these markers namely, Interleukin 17A (IL-17A), Interleukin 10 (IL-10), GM-CSF (CSF2), G-CSF (CSF3), tumor necrosis factor receptor 2 (TNF-RII), tumor necrosis factor beta (TNF-β), TNF-α, Interleukin 1 beta (IL-1β), and Interleukin 1 alpha (IL-1α), were upregulated indicating that the candidate Nanoligomers may be effective in reversing radiation induced immune dysregulation and, therefore, should be tested further. Analysis by clustering using the String Database (StringDB)^49^ indicates the interconnected nature of both the targets and their significantly altered cytokines (Fig.4C). This network has significantly more interactions than expected demonstrating the power of perturbing a member of this gene network (*p*<1.0^-16^). The four clusters from this network: 1. EPO, 2. G-CSF (CSF3), GM-CSF (CSF2), and IL-1A, 3. TNF-α, TNF-β, TNF-RII, and 4. IL-17a, IL-10, and IL-1B indicate that 8U.1_EPO could make for the more interesting case study in animals due to it being the least clustered while still being an effective modulator of cytokine expression.

**Fig.4.**
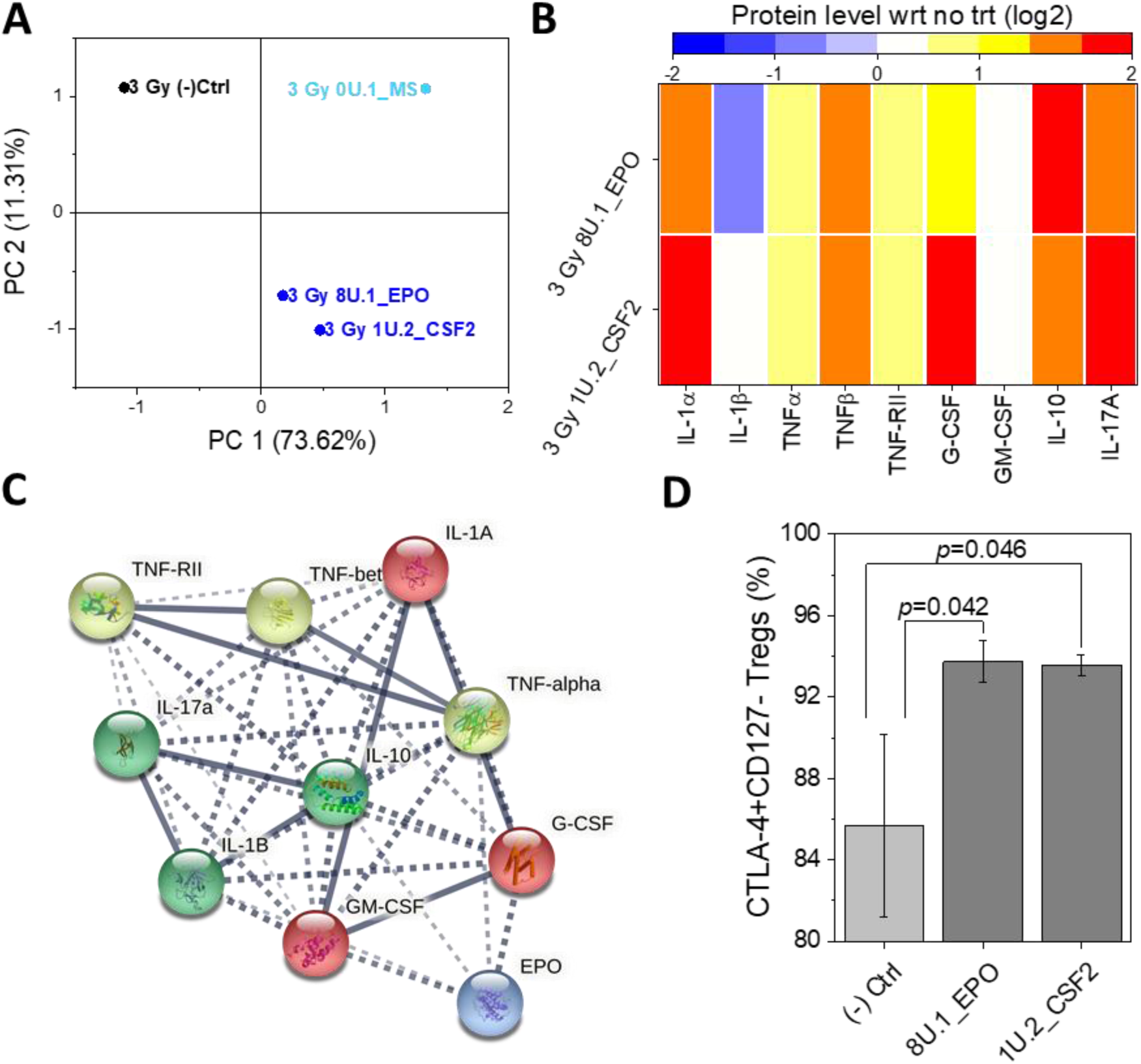
Nanoligomers targeting upregulation of *epo* show favorable immune profile for reversal of immune dysfunction from radiation in human PBMC. A) PBMCs were treated with 10 µM respective Nanoligomer for 24 h followed by total radiation dose of 3 Gy. Principal component analysis of normalized cytokine level shows distinction between targeted Nanoligomers to CSF2 and EPO compared to no treatment and a missense control. B) Nine cytokines shown to be important indicators of radiation induced immune suppression (NASA Twin Study citation) were upregulated by treatment with both 1U.2_CSF2 and 8U.1_EPO. C) The significant cytokines from 1U.2_CSF2 (top) or 8U.1_EPO (bottom) were interrogated and clustered using StringDB to elucidate network conditions and demonstrate the power of using gene expression to modify immune interactions. D) Flow Cytometry of Treg populations show significant increase in CTLA4+CD127- Treg population upon treatment with 8U.1_EPO and 1U.2_CSF2 activators compared to no treatment control.

While approved therapies for H-ARS include Leukine (Sargramostim), the impact of the inflammatory cytokine CSF2 recombinant protein on inducing further immune dysfunction especially on autoimmune diseases, including rheumatoid arthritis and multiple sclerosis, has been implicated.^50^ Its potential role in astrocytosis and microgliosis leading to neurodegenerative diseases has also been revealed, leading to its use as a countermeasure target to treat radiation-induced neuropathy.^50–53^ However, the proposed gene-targeting to upregulate *csf2* and *epo* resulted in upregulation of anti-inflammatory cytokines (for example IL-10) and other regulatory chemokines (see Fig.S10 for pathway analysis), pointing to controlled reversal of immune dysfunction, rather than overall increase of inflammatory markers. This also highlights the difference in immune response with recombinant protein and gene therapy targeting protein upregulation.

Regulatory T (Treg) cells are essential for maintaining peripheral tolerance, preventing autoimmunity, and limiting chronic inflammatory diseases. To enumerate Treg cells, PBMCs were collected and stained for flow cytometry (Fig.5A, S11). Here we consider Tregs of relevance as the proportion of T cells that are Interleukin-2 receptor alpha chain positive (CD25+), forkhead box P3, also known as scurfin, positive (FoxP3+), and their subpopulation that is Interleukin-7 receptor alpha chain negative (CD127-) and cytotoxic T-lymphocyte-associated protein 4 positive (CTLA4+)^54–56^. To probe the impact of lead targets on Treg cell population, we conducted population-level immune regulation analysis with *csf2* and *epo* upregulating Nanoligomers. To enumerate Treg cells, PBMCs were treated with 8U.1_EPO or 1U.2_CSF2 for 24 hours and then collected and stained for flow cytometry (Fig.4D, S11). We demonstrate that treatment with either 8U.1_EPO or 1U.2_CSF2 causes significant increase in Treg population compared to negative control (phosphate buffered saline (PBS) treated). This observed Treg population increase highlights an additional difference in recombinant protein versus reversible gene therapy, and the potential role in regulating radiation dysfunction in multiple protein expression pathways using gene regulation networks.

**Fig 5.**
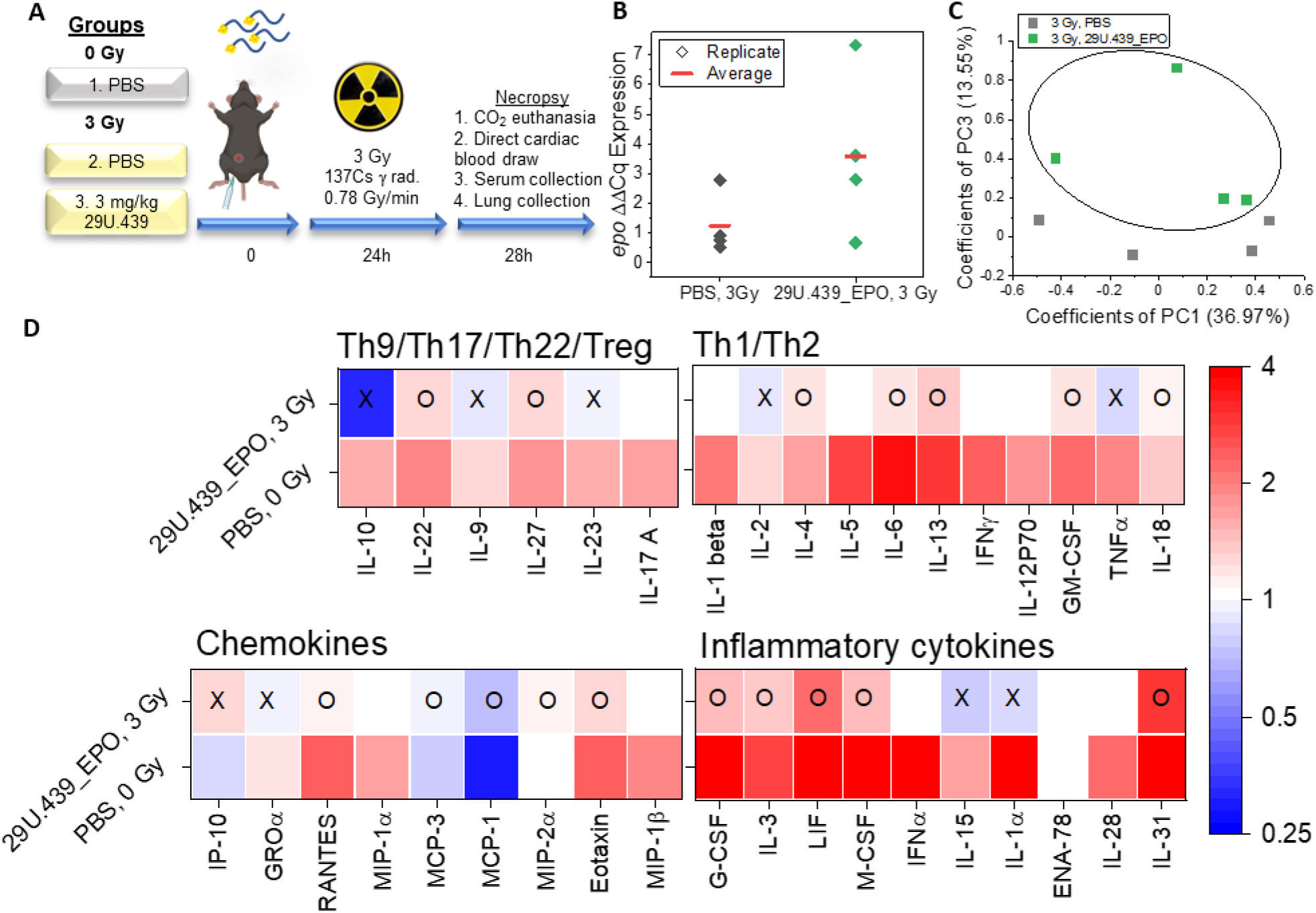
Reversal of immunosuppression in murine radiation model. **A)** Three groups of four female C57BL/6NCrl mice were obtained from Charles River Laboratories at 6 weeks of age and allowed to acclimate for 2 weeks prior to being irradiated. At time 0, respective treatments via intraperitoneal injection were administered to each group as follows: Group 1, PBS, Group 2, PBS, and Group 3, 3 mg/kg 29U.439_EPO. At 24 hours post injection, mice in groups 2 and 3 were given whole-body irradiation using 137Cs γ-rays radiation at a rate of 0.78 Gy/min until total dose of 3 Gy. At 28 hours post injection (4 hours radiation recovery), all groups were euthanized by carbon dioxide and tissue collection was conducted. **B)** RT-qPCR analysis of murine lung tissues from group 2 and group 3 showed increased expression of EPO with treatment by 29U.439_EPO. **C-D)** Lung tissues from all three groups were analyzed for cytokine levels by 36-plex ProcartaPlex assay. The cytokine levels were analyzed by principal component analysis (**C**), and we observed a distinction between irradiated 29U.439_EPO treated animals and untreated irradiated mice in principle component 3 which is a significant component consisting of 13.55% of data variability. (**D)**The 36 cytokines were split into four groups, Th9/Th17/Th22/Treg, Th1/Th2, chemokines, and inflammatory cytokines. After measuring the tissue cytokine levels, we compared group 1 (no treatment, no radiation) and group 3 (treatment with 29U.439_EPO, 3 Gy radiation) against group 2 (no treatment, 3 Gy radiation) to evaluate the reversal of immune suppression by 29U.439_EPO. Heat maps show the fold change in cytokine abundance with respect to group 2 where an “O” indicates 29U.439_EPO changed the cytokine level towards the pre-radiation level and “X” indicates that 29U.439_EPO did not change the cytokine in direction of pre-radiation. Markedly, 18/36 cytokines were altered in the direction of pre-radiation level.

### *In vivo* Testing of Nanoligomer in a Mouse Immunosuppressive Radiation Model

To test *epo* upregulation in a murine model, we used the Nanoligomer bioinformatics tool and synthesis pipeline (Fig.1) and produced 29U.439_EPO. With an existing murine radiation model,^11, 57–64^ 29U.439_EPO was tested as a radioprotector/radiomitigator agent in a pilot study for reversal of radiation-induced immune dysfunction in lung tissue (Fig.5A). Three groups of 4 female C57BL/6NCrl mice were administered either vehicle negative control (group 1 and group 2) or 3 mg/kg 29U.439_EPO (group 3) via intraperitoneal (IP) injection. In previous studies, IP injection has shown high bioavailability of Nanoligomer in lung tissue over 24 hours (data not shown). Twenty-four hours post injection, two groups (group 2 (vehicle) and group 3 (29U.439_EPO)) were irradiated to the whole body with 3 Gy of ^137^Cs γ-rays. All three groups were euthanized 4 hours after irradiation. No animals showed signs of distress during the treatment with Nanoligomers. To evaluate the effect of treatment, blood serum and lung tissue (for carcinogenesis proxy) were harvested.^65^ Blood serum was used to evaluate systemic immune response to Nanoligomer. Based on four key markers, IL-1β, Interleukin 6 (IL-6), interferon gamma (IFN-γ), and TNF-α, we observed no statistically significant increase in systemic inflammatory response to nanoligomer treatment (Fig.S12). One mouse treated with 29U.439 had slightly elevated TNF-α (2.07 pg/mL) but was an outlier in the group so we did not attribute this increase to Nanoligomer treatment. In all 36 assayed factors, there was no statistical difference in the serum levels of cytokines (Fig.S12). This is in contrast to other antisense oligomer therapies which have shown non-specific inflammation in response to administration of the therapeutic molecules .^66, 67^ Further, rapid biodistribution of Nanoligomers in different organs, no accumulation in first pass organs, and renal clearance (data not shown) also solves many outstanding issues with antisense oligonucleotide delivery and pharmacokinetics.^68–70^

Lung tissue was next processed for 36-plex cytokine and chemokine analysis (Invitrogen Cytokine & Chemokine 36-Plex Mouse ProcartaPlex Panel 1A), and RNA was extracted for real-time quantitative polymerase chain reaction (RT-qPCR) of *epo* (see Methods). As shown in Fig.5B, RT-qPCR of RNA shows 29U.439_EPO caused 2-fold increase in relative gene expression of *epo* compared to no treatment (reference genes *actB* and *rpl32*). This demonstrates that 29U.439_EPO was delivered to the lung tissue and upregulated *epo* expression.

Measurement of cytokines and chemokines in the 36-plex panel validated that the chosen irradiated murine model (Fig.S13-16) closely mimics the immunosuppressive radiation response, as seen in long spaceflight missions^5, 71^ and cancer radiotherapy. Markedly, 18 cytokines and chemokines were significantly downregulated, and one was significantly upregulated (monocyte chemoattractant protein-1 (MCP-1 or CCL2)) in PBS, 3 Gy compared to PBS, 0 Gy. This demonstrates that the immune suppression from 3 Gy irradiation is broad.

We performed principal component analysis on the 3 Gy no treatment and 3 Gy, 29U.439_EPO mouse cytokine panel results. Based on the Scree plot and scores, PC1 and PC3, which account for 36.97% and 13.56% of variability, respectively, demonstrate a distinction between the two groups. This highlights that treatment with Nanoligomer changed the cytokine profile in mouse lung tissue. These changes are spread across many of the 36 cytokines and chemokines measured demonstrating broad alteration of the immune response during radiation.

The exact cytokine profile associated with radiation protection is difficult to evaluate and is not well understood. To evaluate *epo* upregulation by 29U.439_EPO, we compared 3 Gy without treatment to 3 Gy with 29U.439_EPO treatment and determined how treatment alters cytokine levels towards the no radiation, no treatment control as an indication of radioprotection and/or reversal of immune suppression due to radiation. Examination of the individual cytokine levels shows that treatment with 29U.439_EPO caused many cytokine and chemokine levels to trend towards to the PBS, 0 Gy lung levels. 29U.439_EPO, 3 Gy significantly decreased IL-10 expression in the murine lung compared to PBS, 3 Gy but this level was not statistically different from the PBS, 0 Gy group (Fig.S16, *p*=0.125) indicating no significant deviation from basal level of IL-10 (Fig.S12). Seventeen markers were reversed towards no radiation levels: T_H_9/T_H_17/T_H_22/Treg type cytokines Interleukin 22 (IL-22) and Interleukin 27 (IL-27), Chemokines: Regulated upon Activation, Normal T Cell Expressed and Presumably Secreted (RANTES), Monocyte Chemotactic Protein 3 (MCP-3), Monocyte Chemoattractant Protein-1 (MCP-1), Macrophage inflammatory protein-2 alpha (MIP-2α), and Eotaxin, T_h_1/Th2 type cytokines IL-4, IL-6, Interleukin 13 (IL-13), GM-CSF, and IL-18, and Inflammatory Cytokines: G-CSF, IL-3, Leukemia inhibitory factor (LIF), M-CSF, and Interleukin 31 (IL-31).While the importance of regulate the Th1/Th2 cytokines has long been appreciated in maintaining the balance between protection and immunopathology,^72^ many recent studies have shown the vital role circulating Th1, Th2, Th9, Th17, Th22, and Treg levels play and their therapeutic significance.^73, 74^ Therefore, the dysfunction created in these four important immunological responses due to radiation, and their trend towards the non-irradiated state due to our radiation countermeasure supports further investigation and potential of 29U.439_EPO. This study focused on acute radiation; however, the safety profile (data not shown) allows for Nanoligomers to be given as a dose regimen across long term exposure as required in long space missions. Our future studies will focus on 1) higher doses of Nanoligomer to maximize gene upregulation, 2) alternate routes of administration such as intravenous to study the effect on addition tissue targets, and 3) multiplexing of 29U.439_EPO with additional targets identified from our PBMC model of radiation dysregulation.

## Conclusion

We have demonstrated a platform for rapid therapeutic lead identification from complex disease pathways, through regulation and gene perturbation of a number of upstream regulators and canonical pathways. We also showed that immune dysfunction caused by radiation exposure through a network of different protein expression can be reversed through identification of single key gene target (*epo*) identified here. While existing therapeutic modalities such as recombinant proteins can effectively treat specific protein expression dysfunction, the Nanoligomer induced reversible gene expression modulation can have important therapeutic application as it combines specificity and ease of delivery, with the ability to target key gene networks to alleviate dysfunction in a number of key protein expression. This represents both a strong target identification platform as well as a therapeutic modality.

Nanoligomers™ as a therapeutic modality offer several advantages beyond platform-based drug-discovery and a single key upregulator identified above. Briefly, this new modality provides a *reversible* up- and down-regulation of any gene target (both gain and loss of function available through transcriptional and translational regulation) without any immunogenic response and good biodistribution characteristics (facile delivery, high bioavailability, and rapid clearance). This is in contrast with other antisense oligonucleotide methods that result in immunogenic response and accumulation in first pass organs like liver and kidney, and demonstrate only translational gene knock-down, and show lower binding affinity and specificity. These attributes make Nanoligomer therapeutic modality and platform as an important tool in development of next-generation countermeasures to make space travel safer.

## Methods

### Nanoligomer Design

Gene targets were determined, and their DNA and transcript sequences used as input to our bioinformatics tool. In one embodiment of our ranking algorithm, we used Naïve-Bayes classifiers^37–42^ to utilize potential match/mismatch of each candidate along the human genome (*C_iMM_*, coefficients indicating number of mismatch across human genome along with specific nucleotide identity A-A, A-C, A-G, C-A, etc., Fig.1B, Supplementary Table S1). To calculate a cumulative/combined score for the ranking based on binding specificity, we assign weighing coefficient (*ω_iMM_*) as the ratio of experimentally measured dissociation constant (*K_D_*) with respective mismatches 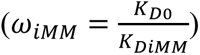. Taking product of the coefficient of mismatch (*C_iMM_*) and the respective weighing coefficient provides the scoring based on specificity. To account for binding affinity of candidates, we used their respective estimated melting temperatures (*T_M_*) to obtain coefficient *ω_TM_* as 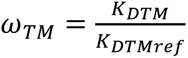, where dissociation constant can be estimated from melting temperatures using the thermodynamic relationship 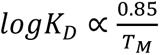 and calculated as 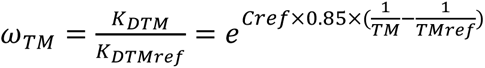. Combining the effects of binding specificity and binding affinity provides a 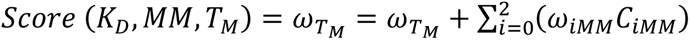. Here 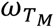 is the weighing coefficient for binding energy, *ω*_*iMM*_ is the weight coefficient for target specificity (respective off-targets), and *C*_*iMM*_ is the number of respective matches throughout the human genome.

### Nanoligomer Synthesis

Nanoligomers™ were synthesized by Sachi Bioworks using patented method. Briefly, up to six different modules based on peptide sequences were synthesized using high-throughput solid phase peptide synthesis. They were purified and assembled into Nanoligomer using the affinity module and coated gold nanoparticle formulation. Following the assembly of Nanoligomer components, molecules were processed using size exclusion centrifugation filters. Purified Nanoligomer concentration was determined using ultraviolet-visible spectroscopy (Thermo Scientific NanoDrop) and known optical parameters. Final Nanoligomer product was filter sterilized prior to use in experiments. In all experiments, Nanoligomer were added directly to cell culture or administered to mouse in buffer without addition of additional delivery reagents. For all studies, a vehicle control was given to no treatment populations.

### Peripheral blood mononuclear cell (PBMC) culture

Commercially prepared PBMCs were purchased from ZenBio and stored in liquid nitrogen until use. Cells were recovered from freeze using manufacturer protocol as follows. A vial of PBMC was rapidly thawed in a 37 ℃ water bath until just prior to complete thawing. The cells were mixed by pipetting and aseptically transferred to a 50 ml conical tube that contained 1 mL prewarmed Lymphocyte Medium (LM). The PBMC vial was rinsed with an additional 1 mL prewarmed LM and combined in the 50 mL conical. A remainder of prewarmed LM was added drop wise to the 50 mL conical until reaching a volume of 25 mL. Cells were centrifuged at 400 time gravity (xg) at room temperature for 10 minutes. The supernatant was removed leaving the cell pellet which was resuspended in LM and cells were enumerated by hemocytometer and Trypan Blue counting. Cells were resuspended to desired density in complete Roswell Park Memorial Institute (RPMI) media prepared with RPMI-1640 with Glutamax (Gibco 72400) and addition of 10% heat inactivated fetal bovine serum and 1% Penicillin/Streptomycin.

### Nanoligomer treatment of PBMC

PBMCs were prepared fresh as described from freeze. For Nanoligomer treatment microplate assays, 500,000 PBMCs were seeded into a 96 well plate at a volume of 200 µL per well. PBMCs were treated with respective Nanoligomer at 10 µM or vehicle negative control. After 24 hours of treatment, cells were exposed to respective radiation treatment and allowed to recover for 4 hours after which the cells were processed for downstream assays.

### Cytokine quantification in PBMC

PBMC culture supernatants were analyzed for secreted proteins with Immune Monitoring 65- Plex Human ProcartaPlex Panel (Thermo Fisher Scientific, Carlsbad, CA) as per manufacturer’s instructions (all proteins assayed shown in Table S5). The assay was read on a Luminex MAGPIX xMAP instrument. Quantification was performed in xPONENT Luminex software where standard curves were created using eight, four-fold dilutions of assay protein standards.

### RNA expression quantification in PBMC

RNA expression in PBMC was quantified using Invitrogen QuantiGene plex gene expression assay. For all gene measurements, three reference genes (Glyceraldehyde-3-Phosphate Dehydrogenase (GAPDH), Phosphoglycerate Kinase 1 (PGK1), and Glucuronidase Beta (GUSB)) were included in addition to the target gene. PBMC were treated with Nanoligomers as described above. Following treatment, PBMCs were pelleted in the round bottom 96-well culture plate at 400 xg for 10 min. Media was removed and cells were lysed following manufacturers instruction. Briefly, lysis mixture was prewarmed at 37 ℃, Proteinase K was added, and the lysis mixture was diluted to a working solution with 2 volumes of nuclease-free water. Cells were lysed at 1000 cells/µL by pipetting up and down 15 times, vortexing at max speed for 1 min, and incubating at 50-55 ℃ for 30 min. After incubation, cells were vortexed in 1 min intervals until complete lysis was confirmed by microscopy. The resultant solutions were processed in the QuantiGene assay and analyzed on the Luminex MAGPIX xMAP instrument.

### RNA expression quantification in mouse tissues

Tissues were stored at -80 ℃ in AllProtect (QIAgen) and removed immediately prior to processing. Briefly, tissues were homogenized with beads at a frequency of 30 Hz for a total of two minutes (four 30 s bursts) using Tissue Lyser II system (Thermo Fisher). RNA was extracted using a miRNeasy Micro Kit (QIAgen, USA) per manufacturer protocol. RNA (500 ng) was reverse transcribed to complementary DNA (cDNA) using High-Capacity cDNA Reverse Transcription Kit (Applied Biosystems). Quantitative real-time polymerase chain reaction (RT-qPCR) was performed using Fast SYBR Green Master Mix (Applied Biosystems) and custom intron-spanning primers (Integrated DNA Technologies, Inc.) shown in Table S4 on Bio-Rad CFX96 Touch Real-Time PCR Detection System. Gene expression for each gene of interest was analyzed following the delta-delta cycle threshold (Ct) method relative to the reference genes, ribosomal protein L32 (RPL32) and beta actin (ACTB).

### Murine model study

All animal work was approved by the Institutional Animal Care and Use Committee at Colorado State University under protocol 1891. The animal care facilities at Colorado State University are AAALAC accredited. Female C57BL/6NCrl mice were obtained from Charles River Laboratories at 6 weeks of age and allowed to acclimate for 2 weeks prior to being irradiated. Mice were ear punched for identification one week prior to experiment to allow for healing and return to immune homeostasis. Nanoligomer (Group 3) or phosphate buffered saline (vehicle negative control, Group 1 and 2) were administered at day/time 0. Nanoligomer was given at 3 mg/kg (see supplemental Fig.S12). At time 24 hours post injection, unanesthetized mice were exposed to 3 Gy whole-body irradiation with 137Cs gamma-rays radiation at a dose rate of 0.78 Gy/minute. At time 28 hours (4 hours recovery after radiation), animals were euthanized by carbon dioxide and blood was immediately drawn by direct cardiac puncture draw. Whole blood was kept once ice for 10 min to clot, following by centrifugation at 2000 xg for 10 min, and serum was removed by pipette, transferred to a fresh tube, and frozen. Lung tissues were then collected, and samples were immediately flash frozen or submerged in AllProtect (Qiagen) (first at room temperature for stabilization) and stored at -80 ℃ until processing.

### Multiplex cytokine panel

Flash frozen tissues were manually homogenized using a mortar and pestle and syringe disruption until a homogenate solution was formed in 500 µL /100mg tissue cell lysis buffer (EPX-99999-000). Homogenate was centrifuged at 16,000 xg for 10 min at 4 ℃ and the supernatant was saved in a new tube as the sample for analysis. Protein content of the sample was determined using Bio-Rad’s DC Protein Assay Kit with BSA standard and read using a Tecan GENios microplate reader. All samples were then diluted to 10 mg protein/mL with PBS. Quantification of cytokines was performed using 25 µL of protein or serum in Invitrogen Cytokine & Chemokine Convenience 36-Plex Mouse ProcartaPlex Panel 1A (EPXR360-26092-901) (all proteins assayed shown in Table S5) following manufacturers instruction and was read on a Luminex MAGPIX xMAP instrument. Quantification was done using xPONENT Luminex software where standard curves were created using eight, four-fold dilutions of assay protein standards.

### T-regulatory cells (Tregs) Enumeration

T-regulatory cells (Tregs) are a subset of cluster of differentiation 4 positive (CD4+) T cells that are CD25+FoxP3+ and their subpopulation that is CD127-CTLA4+. To enumerate Tregs following Nanoligomer treatment, PBMCs were collected, washed with staining buffer (PBS with 2% bovine serum albumin and 2 mM EDTA), and stained for flow cytometry. The following antibodies were used for surface staining: BV785-labelled anti-CD3 antibody, PerCP/Cy5.5-labelled anti-CD4, Alexa405-labelled anti-CD8 antibody, APC-Cy7-labeled anti-CD25, and FITC-labeled anti-CD127. Intracellular staining for PE-labeled anti-FoxP3 antibody, APC-labeled anti-CTLA-4 antibody (BD Biosciences) was performed using the FoxP3 staining buffer set. Flow cytometry analysis was done using LSR II (BD Biosciences). Isotype controls and/or FMOs (Fluorescence Minus One) were used to determine gating boundaries. Data analyzed using FlowJo software (FlowJo, LLC).

### Data analysis

Error bars represent two standard deviations of biological replicates. In all cases, significance designated with an asterisk (*) is defined as *p* < 0.05 for a 95% confidence interval. Clustering and dendrogram as well as principal component analysis were performed Origin.

## Funding Sources

We acknowledge financial support from National Aeronautics and Space Administration SBIR Phase I Contract 80NSSC21C0242 to Sachi Bioworks, funding from Sachi Bioworks, and funding from NASA grant NNX15AK13G to MMW.

## Acknowledgements

The authors sincerely thank Dr. Charles Preston Neff - Technical Director and Garrett Hedlund from Flow Cytometry Facility, University of Colorado Anschutz, for their help with flow cytometry analysis. The authors also thank Prof. Alexander Brandl-Director, Irradiation Services Laboratory and Justin Bell at Colorado State University for their time spent in coordination and irradiation of PBMC samples.

## Author Contributions

The manuscript was written through contributions of all authors. C.M.C., S.S., P.N., and A.C. designed all experiments and conducted Nanoligomer design, synthesis, *in vitro* testing, and data analysis/preparation. C.M.C., C.F., M.M.W, and P.N. conducted *in vivo* studies. All authors discussed the results and edited the manuscript.

## Competing Interests

A.C., P.N., S.S., and C.M.C. are employed by the for-profit company Sachi Bioworks where this technology was developed. A.C. and P.N. are the founders of Sachi Bioworks, and P.N. has filed a patent on this technology.

## References

(1) Cucinotta, F. A.; Hu, S.; Schwadron, N. A.; Kozarev, K.; Townsend, L. W.; Kim, M. H. Y. Space Radiation Risk Limits and Earth-Moon-Mars Environmental Models. *Sp*. Weather 2010, 8 (12), 1–12. https://doi.org/10.1029/2010SW000572.

(2) Stanishevsky, A.; Nagaraj, B.; Melngailis, J.; Ramesh, R.; Khriachtchev, L.; McDaniel, E. Radiation Damage and Its Recovery in Focused Ion Beam Fabricated Ferroelectric Capacitors. J. Appl. Phys. 2002, 92 (6), 3275–3278. https://doi.org/10.1063/1.1489069.

(3) Singh, V. K.; Newman, V. L.; Seed, T. M. Colony-Stimulating Factors for the Treatment of the Hematopoietic Component of the Acute Radiation Syndrome (H-ARS): A Review. Cytokine 2015, 71 (1), 22–37. https://doi.org/10.1016/j.cyto.2014.08.003.

(4) Crucian, B. E.; Choukèr, A.; Simpson, R. J.; Mehta, S.; Marshall, G.; Smith, S. M.; Zwart, S. R.; Heer, M.; Ponomarev, S.; Whitmire, A.; Frippiat, J. P.; Douglas, G.; Lorenzi, H.; Buchheim, J. I.; Makedonas, G.; Ginsburg, G. S.; Mark Ott, C.; Pierson, D. L.; Krieger, S. S.; Baecker, N.; Sams, C. Immune System Dysregulation during Spaceflight: Potential Countermeasures for Deep Space Exploration Missions. Frontiers in Immunology. 2018. https://doi.org/10.3389/fimmu.2018.01437.

(5) Garrett-Bakelman, F. E.; Darshi, M.; Green, S. J.; Gur, R. C.; Lin, L.; Macias, B. R.; McKenna, M. J.; Meydan, C.; Mishra, T.; Nasrini, J.; Piening, B. D.; Rizzardi, L. F.; Sharma, K.; Siamwala, J. H.; Taylor, L.; Vitaterna, M. H.; Afkarian, M.; Afshinnekoo, E.; Ahadi, S.; Ambati, A.; Arya, M.; Bezdan, D.; Callahan, C. M.; Chen, S.; Choi, A. M. K.; Chlipala, G. E.; Contrepois, K.; Covington, M.; Crucian, B. E.; De Vivo, I.; Dinges, D. F.; Ebert, D. J.; Feinberg, J. I.; Gandara, J. A.; George, K. A.; Goutsias, J.; Grills, G. S.; Hargens, A. R.; Heer, M.; Hillary, R. P.; Hoofnagle, A. N.; Hook, V. Y. H.; Jenkinson, G.; Jiang, P.; Keshavarzian, A.; Laurie, S. S.; Lee-McMullen, B.; Lumpkins, S. B.; MacKay, M.; Maienschein-Cline, M. G.; Melnick, A. M.; Moore, T. M.; Nakahira, K.; Patel, H. H.; Pietrzyk, R.; Rao, V.; Saito, R.; Salins, D. N.; Schilling, J. M.; Sears, D. D.; Sheridan, C. K.; Stenger, M. B.; Tryggvadottir, R.; Urban, A. E.; Vaisar, T.; Van Espen, B.; Zhang, J.; Ziegler, M. G.; Zwart, S. R.; Charles, J. B.; Kundrot, C. E.; Scott, G. B. I.; Bailey, S. M.; Basner, M.; Feinberg, A. P.; Lee, S. M. C.; Mason, C. E.; Mignot, E.; Rana, B. K.; Smith, S. M.; Snyder, M. P.; Turek, F. W. The NASA Twins Study: A Multidimensional Analysis of a Year-Long Human Spaceflight. Science (80-.). 2019. https://doi.org/10.1126/science.aau8650.

(6) Guéguinou, N.; Huin-Schohn, C.; Bascove, M.; Bueb, J.-L.; Tschirhart, E.; Legrand-Frossi, C.; Frippiat, J.-P. Could Spaceflight-Associated Immune System Weakening Preclude the Expansion of Human Presence beyond Earth’s Orbit? J. Leukoc. Biol. 2009. https://doi.org/10.1189/jlb.0309167.

(7) Makedonas, G.; Mehta, S.; Choukèr, A.; Simpson, R. J.; Marshall, G.; Orange, J. S.; Aunon-Chancellor, S.; Smith, S. M.; Zwart, S. R.; Stowe, R. P.; Heer, M.; Ponomarev, S.; Whitmire, A.; Frippiat, J. P.; Douglas, G. L.; Krieger, S. S.; Lorenzi, H.; Buchheim, J. I.; Ginsburg, G. S.; Ott, C. M.; Downs, M.; Pierson, D.; Baecker, N.; Sams, C.; Crucian, B. Specific Immunologic Countermeasure Protocol for Deep-Space Exploration Missions. Front. Immunol. 2019. https://doi.org/10.3389/fimmu.2019.02407.

(8) Vuong, C.; Saenz, H. L.; Götz, F.; Otto, M. Reactivation and Shedding of Cytomegalovirus in Astronauts during Spaceflight. J. Infect. Dis. 2000. https://doi.org/10.1086/317624.

(9) Singh, V. K.; Romaine, P. L. P.; Seed, T. M. Medical Countermeasures for Radiation Exposure and Related Injuries: Characterization of Medicines, FDA-Approval Status and Inclusion into the Strategic National Stockpile. Health Physics. 2015. https://doi.org/10.1097/HP.0000000000000279.

(10) Kaur, P.; Asea, A. Radiation-Induced Effects and the Immune System in Cancer. Front. Oncol. 2012, 2. https://doi.org/10.3389/fonc.2012.00191.

(11) Schaue, D.; Kachikwu, E. L.; McBride, W. H. Cytokines in Radiobiological Responses: A Review. Radiation Research. 2012. https://doi.org/10.1667/RR3031.1.

(12) Hérodin, F.; Bourin, P.; Mayol, J. F.; Lataillade, J. J.; Drouet, M. Short-Term Injection of Antiapoptotic Cytokine Combinations Soon after Lethal γ-Irradiation Promotes Survival. Blood 2003. https://doi.org/10.1182/blood-2002-06-1634.

(13) Laiakis, E. C.; Baulch, J. E.; Morgan, W. F. Cytokine and Chemokine Responses after Exposure to Ionizing Radiation: Implications for the Astronauts. Adv. Sp. Res. 2007. https://doi.org/10.1016/j.asr.2006.11.010.

(14) Lierova, A.; Jelicova, M.; Nemcova, M.; Proksova, M.; Pejchal, J.; Zarybnicka, L.; Sinkorova, Z. Cytokines and Radiation-Induced Pulmonary Injuries. Journal of Radiation Research. 2018. https://doi.org/10.1093/jrr/rry067.

(15) Gale, R. P.; Armitage, J. O. Use of Molecularly-Cloned Haematopoietic Growth Factors in Persons Exposed to Acute High-Dose, High-Dose Rate Whole-Body Ionizing Radiations. Blood Reviews. 2021. https://doi.org/10.1016/j.blre.2020.100690.

(16) Reeves, G. Overview of Use of G-CSF and GM-CSF in the Treatment of Acute Radiation Injury. In Health Physics; 2014; Vol. 106. https://doi.org/10.1097/HP.0000000000000090.

(17) Ruef, C.; Coleman, D. L. [GM-CSF and G-CSF: cytokines in clinical application]. Schweiz. Med. Wochenschr. 1991, 121 (12), 397– 412.

(18) Belenkov, A. I.; Shenouda, G.; Rizhevskaya, E.; Cournoyer, D.; Belzile, J.-P.; Souhami, L.; Devic, S.; Chow, T. Y. K. Erythropoietin Induces Cancer Cell Resistance to Ionizing Radiation and to Cisplatin. Mol. Cancer Ther. 2004, 3 (12), 1525–1532. https://doi.org/10.1016/s0167-8140(97)00186-2.

(19) Jin Soo Lee. The Use of Erythropoietin in Radiation Oncology. Cancer Control. 1998, pp 33–40. https://doi.org/10.1177/107327489800502s07.

(20) Neta, R.; Oppenheim, J. J.; Douches, S. D. Interdependence of the Radioprotective Effects of Human Recombinant Interleukin 1 Alpha, Tumor Necrosis Factor Alpha, Granulocyte Colony-Stimulating Factor, and Murine Recombinant Granulocyte-Macrophage Colony-Stimulating Factor. J. Immunol. 1988.

(21) Neta, R.; Oppenheim, J. J.; Schreiber, R. D.; Chizzonite, R.; Ledney, G. D.; MacVittie, T. J. Role of Cytokines (Interleukin 1, Tumor Necrosis Factor, and Transforming Growth Factor β) in Natural and Lipopolysaccharide-Enhanced Radioresistance. J. Exp. Med. 1991. https://doi.org/10.1084/jem.173.5.1177.

(22) Weiss, J. F.; Kumar, K. S.; Walden, T. L.; Neta, R.; Landauer, M. R.; Clark, E. P. Advances in Radioprotection through the Use of Combined Agent Regimens. Int. J. Radiat. Biol. 1990. https://doi.org/10.1080/09553009014550881.

(23) Weiss, J. F.; Landauer, M. R. Radioprotection by Antioxidants. In Annals of the New York Academy of Sciences; 2000. https://doi.org/10.1111/j.1749-6632.2000.tb06175.x.

(24) Michnik, A.; Polaczek-Grelik, K.; Leniak, P.; Drzazga, Z. Effects of Low-Dose Ionizing Radiation on α,β-Globulins Solutions Studied by DSC. J. Therm. Anal. Calorim. 2013, 111 (3), 1845–1852. https://doi.org/10.1007/s10973-012-2687-6.

(25) Birke, G.; Jacobsson, F.; Liljedahl, S. O.; Plantin, L. O.; Wetterfors, J. Catabolism of Albumin and Gamma Globulin after Treatment with Ionising Radiation to the Abdomen. Acta radiol. 1967, 6 (2), 113–121. https://doi.org/10.3109/02841856709138570.

(26) Lima, C. V.; Campos, T. P. R. Kinetics of the Expressions of Radiation-Induced Plasma Proteins of the Cardiac Territory in Electrophoresis. J. Bras. Patol. e Med. Lab. 2016, 52 (3), 171–177. https://doi.org/10.5935/1676-2444.20160029.

(27) Ghandhi, S. A.; Smilenov, L. B.; Elliston, C. D.; Chowdhury, M.; Amundson, S. A. Radiation Dose-Rate Effects on Gene Expression for Human Biodosimetry. BMC Med. Genomics 2015, 8 (1), 22. https://doi.org/10.1186/s12920-015-0097-x.

(28) Smirnov, D. A.; Morley, M.; Shin, E.; Spielman, R. S.; Cheung, V. G. Genetic Analysis of Radiation-Induced Changes in Human Gene Expression. Nature 2009, 459 (7246), 587–591. https://doi.org/10.1038/nature07940.

(29) Azria, D.; Larbouret, C.; Garambois, V.; Kramar, A.; Martineau, P.; Robert, B.; Aillères, N.; Ychou, M.; Dubois, J. B.; Pèlegrin, A. Potentiation of Ionising Radiation by Targeting Tumour Necrosis Factor Alpha Using a Bispecific Antibody in Human Pancreatic Cancer. Br. J. Cancer 2003. https://doi.org/10.1038/sj.bjc.6601362.

(30) Chatterjee, M.; Chakraborty, T.; Tassone, P. Multiple Myeloma: Monoclonal Antibodies-Based Immunotherapeutic Strategies and Targeted Radiotherapy. Eur. J. Cancer 2006. https://doi.org/10.1016/j.ejca.2006.02.016.

(31) Rodriguez-Palacios, A.; Harding, A.; Menghini, P.; Himmelman, C.; Retuerto, M.; Nickerson, K. P.; Lam, M.; Croniger, C. M.; McLean, M. H.; Durum, S. K.; Pizarro, T. T.; Ghannoum, M. A.; Ilic, S.; McDonald, C.; Cominelli, F. The Artificial Sweetener Splenda Promotes Gut Proteobacteria, Dysbiosis, and Myeloperoxidase Reactivity in Crohn’s Disease-Like Ileitis. Inflamm. Bowel Dis. 2018. https://doi.org/10.1093/ibd/izy060.

(32) Nielsen, P. E.; Egholm, M.; Berg, R. H.; Buchardt, O. Sequence-Selective Recognition of DNA by Strand Displacement with a Thymine-Substituted Polyamide. Science (80-.). 1991, 254 (5037), 1497–1500. https://doi.org/10.1126/science.1962210.

(33) Hyrup, B.; Nielsen, P. E. Peptide Nucleic Acids (PNA): Synthesis, Properties and Potential Applications. Bioorg. Med. Chem. 1996, 4 (1), 5–23.

(34) Demidov, V. V.; Potaman, V. N.; Frank-Kamenetskil, M. D.; Egholm, M.; Buchard, O.; Sönnichsen, S. H.; Nlelsen, P. E. Stability of Peptide Nucleic Acids in Human Serum and Cellular Extracts. Biochem. Pharmacol. 1994, 48 (6), 1310–1313. https://doi.org/10.1016/0006-2952(94)90171-6.

(35) Zhao, C.; Hoppe, T.; Setty, M. K. H. G.; Murray, D.; Chun, T.-W.; Hewlett, I.; Appella, D. H. Quantification of Plasma HIV RNA Using Chemically Engineered Peptide Nucleic Acids. Nat. Commun. 2014, 5, 5079. https://doi.org/10.1038/ncomms6079.

(36) Bennett, C. F.; Swayze, E. E. RNA Targeting Therapeutics: Molecular Mechanisms of Antisense Oligonucleotides as a Therapeutic Platform. Annu. Rev. Pharmacol. Toxicol. 2010, 50, 259–293. https://doi.org/10.1146/annurev.pharmtox.010909.105654.

(37) Korshoj, L. E.; Nagpal, P. BOCS: DNA k-Mer Content and Scoring for Rapid Genetic Biomarker Identification at Low Coverage. Comput. Biol. Med. 2019, 110, 196–206. https://doi.org/10.1016/j.compbiomed.2019.05.022.

(38) Korshoj, L. E.; Afsari, S.; Chatterjee, A.; Nagpal, P. Conformational Smear Characterization and Binning of Single-Molecule Conductance Measurements for Enhanced Molecular Recognition. J. Am. Chem. Soc. 2017, 139 (43), 15420–15428. https://doi.org/10.1021/jacs.7b08246.

(39) Afsari, S.; Korshoj, L. E.; Abel, G. R.; Khan, S.; Chatterjee, A.; Nagpal, P.; Abel Jr., G. R.; Khan, S.; Chatterjee, A.; Nagpal, P. Quantum Point Contact Single-Nucleotide Conductance for DNA and RNA Sequence Identification. ACS Nano 2017, 11 (11), 11169– 11181. https://doi.org/10.1021/acsnano.7b05500.

(40) Korshoj, L. E.; Nagpal, P. Diagnostic Optical Sequencing. ACS Appl. Mater. Interfaces 2019, 11 (39), 35587–35596. https://doi.org/10.1021/acsami.9b12568.

(41) Abel, G. R.; Korshoj, L. E.; Otoupal, P. B.; Khan, S.; Chatterjee, A.; Nagpal, P. Nucleotide and Structural Label Identification in Single RNA Molecules with Quantum Tunneling Spectroscopy. Chem. Sci. 2019, 10 (4), 1052–1063. https://doi.org/10.1039/c8sc03354d.

(42) Korshoj, L. E.; Afsari, S.; Khan, S.; Chatterjee, A.; Nagpal, P. Single Nucleobase Identification Using Biophysical Signatures from Nanoelectronic Quantum Tunneling. Small 2017, 13 (11). https://doi.org/10.1002/smll.201603033.

(43) Sen, P.; Kemppainen, E.; Orešič, M. Perspectives on Systems Modeling of Human Peripheral Blood Mononuclear Cells. Frontiers in Molecular Biosciences. 2018. https://doi.org/10.3389/fmolb.2017.00096.

(44) Dobrovolskaia, M. A.; Afonin, K. A. Use of Human Peripheral Blood Mononuclear Cells to Define Immunological Properties of Nucleic Acid Nanoparticles. Nat. Protoc. 2020, 15 (11). https://doi.org/10.1038/s41596-020-0393-6.

(45) Schroecksnadel, K.; Winkler, C.; Wirleitner, B.; Schennach, H.; Fuchs, D. Aspirin Down-Regulates Tryptophan Degradation in Stimulated Human Peripheral Blood Mononuclear Cells in Vitro. Clin. Exp. Immunol. 2005, 140 (1). https://doi.org/10.1111/j.1365-2249.2005.02746.x.

(46) Cheki, M.; Shirazi, A.; Mahmoudzadeh, A.; Bazzaz, J. T.; Hosseinimehr, S. J. The Radioprotective Effect of Metformin against Cytotoxicity and Genotoxicity Induced by Ionizing Radiation in Cultured Human Blood Lymphocytes. Mutat. Res. - Genet. Toxicol. Environ. Mutagen. 2016, 809. https://doi.org/10.1016/j.mrgentox.2016.09.001.

(47) Shimura, T.; Koyama, M.; Aono, D.; Kunugita, N. Epicatechin as a Promising Agent to Countermeasure Radiation Exposure by Mitigating Mitochondrial Damage in Human Fibroblasts and Mouse Hematopoietic Cells. FASEB J. 2019, 33 (6). https://doi.org/10.1096/fj.201802246RR.

(48) Lever, J.; Krzywinski, M.; Altman, N. Principal Component Analysis. Nat. Methods 2017, 14 (7), 641–642. https://doi.org/10.1038/nmeth.4346.

(49) Szklarczyk, D.; Franceschini, A.; Wyder, S.; Forslund, K.; Heller, D.; Huerta-Cepas, J.; Simonovic, M.; Roth, A.; Santos, A.; Tsafou, K. P.; Kuhn, M.; Bork, P.; Jensen, L. J.; Von Mering, C. STRING V10: Protein-Protein Interaction Networks, Integrated over the Tree of Life. Nucleic Acids Res. 2015, 43 (D1). https://doi.org/10.1093/nar/gku1003.

(50) Lotfi, N.; Thome, R.; Rezaei, N.; Zhang, G. X.; Rezaei, A.; Rostami, A.; Esmaeil, N. Roles of GM-CSF in the Pathogenesis of Autoimmune Diseases: An Update. Frontiers in Immunology. 2019. https://doi.org/10.3389/fimmu.2019.01265.

(51) Aram, J.; Francis, A.; Tanasescu, R.; Constantinescu, C. S. Granulocyte-Macrophage Colony-Stimulating Factor as a Therapeutic Target in Multiple Sclerosis. Neurol. Ther. 2019, 8 (1). https://doi.org/10.1007/s40120-018-0120-1.

(52) Dikmen, H. O.; Hemmerich, M.; Lewen, A.; Hollnagel, J. O.; Chausse, B.; Kann, O. GM-CSF Induces Noninflammatory Proliferation of Microglia and Disturbs Electrical Neuronal Network Rhythms in Situ. J. Neuroinflammation 2020, 17 (1). https://doi.org/10.1186/s12974-020-01903-4.

(53) Shiomi, A.; Usui, T.; Mimori, T. GM-CSF as a Therapeutic Target in Autoimmune Diseases. Inflamm. Regen. 2016, 36 (1). https://doi.org/10.1186/s41232-016-0014-5.

(54) Sakaguchi, S.; Miyara, M.; Costantino, C. M.; Hafler, D. A. FOXP3 + Regulatory T Cells in the Human Immune System. Nature Reviews Immunology. 2010. https://doi.org/10.1038/nri2785.

(55) Gavin, M. A.; Torgerson, T. R.; Houston, E.; DeRoos, P.; Ho, W. Y.; Stray-Pedersen, A.; Ocheltree, E. L.; Greenberg, P. D.; Ochs, H. D.; Rudensky, A. Y. Single-Cell Analysis of Normal and FOXP3-Mutant Human T Cells: FOXP3 Expression without Regulatory T Cell Development. Proc. Natl. Acad. Sci. U. S. A. 2006, 103 (17). https://doi.org/10.1073/pnas.0509484103.

(56) Neff, C. P.; Rhodes, M. E.; Arnolds, K. L.; Collins, C. B.; Donnelly, J.; Nusbacher, N.; Jedlicka, P.; Schneider, J. M.; McCarter, M. D.; Shaffer, M.; Mazmanian, S. K.; Palmer, B. E.; Lozupone, C. A. Diverse Intestinal Bacteria Contain Putative Zwitterionic Capsular Polysaccharides with Anti-Inflammatory Properties. Cell Host Microbe 2016. https://doi.org/10.1016/j.chom.2016.09.002.

(57) Ha, C. T.; Li, X. H.; Fu, D.; Moroni, M.; Fisher, C.; Arnott, R.; Srinivasan, V.; Xiao, M. Circulating Interleukin-18 as a Biomarker of Total-Body Radiation Exposure in Mice, Minipigs, and Nonhuman Primates (NHP). PLoS One 2014, 9 (10). https://doi.org/10.1371/journal.pone.0109249.

(58) Ha, C. T.; Li, X.; Fu, D.; Xiao, M. Circulating IL-18 Binding Protein (IL-18BP) and IL-18 as Dual Biomarkers of Total-Body Irradiation in Mice. Radiat. Res. 2016, 185 (4). https://doi.org/10.1667/RR14238.1.

(59) Hill, G. R.; Crawford, J. M.; Cooke, K. R.; Brinson, Y. S.; Pan, L.; Ferrara, J. L. M. Total Body Irradiation and Acute Graft-versus-Host Disease: The Role of Gastrointestinal Damage and Inflammatory Cytokines. Blood 1997, 90 (8). https://doi.org/10.1182/blood.v90.8.3204.

(60) Liu, X.; Liu, Z.; Wang, D.; Han, Y.; Hu, S.; Xie, Y.; Liu, Y.; Zhu, M.; Guan, H.; Gu, Y.; Zhou, P. K. Effects of Low Dose Radiation on Immune Cells Subsets and Cytokines in Mice. Toxicol. Res. (Camb). 2020, 9 (3). https://doi.org/10.1093/TOXRES/TFAA017.

(61) Bogdándi, E. N.; Balogh, A.; Felgyinszki, N.; Szatmári, T.; Persa, E.; Hildebrandt, G.; Sáfrány, G.; Lumniczky, K. Effects of Low-Dose Radiation on the Immune System of Mice after Total-Body Irradiation. Radiat. Res. 2010, 174 (4). https://doi.org/10.1667/RR2160.1.

(62) McKelvey, K. J.; Hudson, A. L.; Back, M.; Eade, T.; Diakos, C. I. Radiation, Inflammation and the Immune Response in Cancer. Mammalian Genome. 2018. https://doi.org/10.1007/s00335-018-9777-0.

(63) Multhoff, G.; Radons, J. Radiation, Inflammation, and Immune Responses in Cancer. Frontiers in Oncology. 2012. https://doi.org/10.3389/fonc.2012.00058.

(64) Pan, S.; Wang, J.; Wu, A.; Guo, Z.; Wang, Z.; Zheng, L.; Dai, Y.; Zhu, L.; Nie, J.; Hei, T. K.; Zhou, G.; Li, Y.; Li, B.; Hu, W. Radiation Exposure–Induced Changes in the Immune Cells and Immune Factors of Mice With or Without Primary Lung Tumor. Dose-Response 2020, 18 (2). https://doi.org/10.1177/1559325820926744.

(65) Shay, J. W.; Cucinotta, F. A.; Sulzman, F. M.; Coleman, C. N.; Minna, J. D. From Mice and Men to Earth and Space: Joint NASA-NCI Workshop on Lung Cancer Risk Resulting from Space and Terrestrial Radiation. In Cancer Research; 2011; Vol. 71. https://doi.org/10.1158/0008-5472.CAN-11-2546.

(66) Meng, Z.; Lu, M. RNA Interference-Induced Innate Immunity, off-Target Effect, or Immune Adjuvant? Front. Immunol. 2017, 8 (MAR), 1–7. https://doi.org/10.3389/fimmu.2017.00331.

(67) Sioud, M. Induction of Inflammatory Cytokines and Interferon Responses by Double-Stranded and Single-Stranded SiRNAs Is Sequence-Dependent and Requires Endosomal Localization. J. Mol. Biol. 2005, 348 (5), 1079–1090. https://doi.org/10.1016/j.jmb.2005.03.013.

(68) Geary, R. S.; Norris, D.; Yu, R.; Bennett, C. F. Pharmacokinetics, Biodistribution and Cell Uptake of Antisense Oligonucleotides. Advanced Drug Delivery Reviews. 2015, pp 46–51. https://doi.org/10.1016/j.addr.2015.01.008.

(69) Thompson, J. D.; Kornbrust, D. J.; Foy, J. W. D.; Solano, E. C. R.; Schneider, D. J.; Feinstein, E.; Molitoris, B. A.; Erlich, S. Toxicological and Pharmacokinetic Properties of Chemically Modified SiRNAs Targeting P53 RNA Following Intravenous Administration. Nucleic Acid Ther. 2012, 22 (4). https://doi.org/10.1089/nat.2012.0371.

(70) Geary, R. S.; Yu, R. Z.; Siwkowski, A.; Levin, A. A. Pharmacokinetic/Pharmacodynamic Properties of Phosphorothioate 2′-O-(2-Methoxyethyl)-Modified Antisense Oligonucleotides in Animals and Man. In Antisense Drug Technology: Principles, Strategies, and Applications, *Second Edition*; 2007. https://doi.org/10.1201/9780849387951.ch11.

(71) da Silveira, W. A.; Fazelinia, H.; Rosenthal, S. B.; Laiakis, E. C.; Kim, M. S.; Meydan, C.; Kidane, Y.; Rathi, K. S.; Smith, S. M.; Stear, B.; Ying, Y.; Zhang, Y.; Foox, J.; Zanello, S.; Crucian, B.; Wang, D.; Nugent, A.; Costa, H. A.; Zwart, S. R.; Schrepfer, S.; Elworth, R. A. L.; Sapoval, N.; Treangen, T.; MacKay, M.; Gokhale, N. S.; Horner, S. M.; Singh, L. N.; Wallace, D. C.; Willey, J. S.; Schisler, J. C.; Meller, R.; McDonald, J. T.; Fisch, K. M.; Hardiman, G.; Taylor, D.; Mason, C. E.; Costes, S. V.; Beheshti, A. Comprehensive Multi-Omics Analysis Reveals Mitochondrial Stress as a Central Biological Hub for Spaceflight Impact. Cell 2020. https://doi.org/10.1016/j.cell.2020.11.002.

(72) Singh, V. K.; Mehrotra, S.; Agarwal, S. S. The Paradigm of Th1 and Th2 Cytokines: Its Relevance to Aotoimmunity and Allergy. Immunologic Research. 1999. https://doi.org/10.1007/bf02786470.

(73) Ye, J.; Wang, Y.; Wang, Z.; Ji, Q.; Huang, Y.; Zeng, T.; Hu, H.; Ye, D.; Wan, J.; Lin, Y. Circulating Th1, Th2, Th9, Th17, Th22, and Treg Levels in Aortic Dissection Patients. Mediators Inflamm. 2018, 2018. https://doi.org/10.1155/2018/5697149.

(74) Cui, G. TH9, TH17, and TH22 Cell Subsets and Their Main Cytokine Products in the Pathogenesis of Colorectal Cancer. Frontiers in Oncology. 2019. https://doi.org/10.3389/fonc.2019.01002.

