## Supplemental Information for "Reversing Radiation-Induced Immunosuppression Using a New Therapeutic Modality"

**TABLE OF CONTENTS:**

**Supplementary Tables S1-4**

**Supplementary Figures S1-17**

**Supplementary References**

**Table S1.** List of gene targets and genome reference sequence ID used for Nanoligomer targeting.

| Gene Name | Reference Sequence |
|---|---|
| Granulocyte-Macrophage Colony-Stimulating Factor (CSF2) | NG_033024.1 |
| Granulocyte Colony-Stimulating Factor (CSF3) | NM_000759.4 |
| Stem Cell Factor (SCF) | NG_012098.2 |
| FMS-like Tyrosine Kinase 3 Ligand (FTL3LG) | NM_001204502.2 |
| Thrombopoietin (THPO) | NG_012136.1 |
| Erythropoietin (EPO) | NG_021471.2 |
| Interleukin-3 (IL-3) | NM_000588.4 |
| Stromal Cell-Derived Factor-1 (SDF-1) | NG_016861.2 |
| Interleukin-1α (IL-1α) | NG_008850.1 |
| Interleukin-1β (IL-1β) | NG_008851.1 |
| Tumor Necrosis Factor-α (TNF-α) | NG_007462.1 |
| Interleukin-6 (IL-6) | NG_011640.1 |

**Table S2.** The upper quartile of cytokines with the greatest impact on respective principal component are shown here. The contributing cytokines were compared to data from the NASA twin study[1] and their significant alteration during inflight vs preflight, postflight vs preflight, and postflight vs. inflight are indicated with an ampersand. N/A indicates the cytokine was not measured in the NASA study. The strong overlap between cytokines with a governing role on principal component analysis and those which were significantly different in the NASA twin study[1] highlights the biological value of using the principal components to select top Nanoligomers.

| Top Quartile PC1 | Significant inflght vs preflight | Significant post-flight vs preflight | Significant post-flight vs inflight | Top Quartile PC2 | Significant inflght vs preflight | Significant post-flight vs preflight | Significant post-flight vs inflight | Top Quartile PC3 | Significant inflght vs preflight | Significant post-flight vs preflight | Significant post-flight vs inflight |
|---|---|---|---|---|---|---|---|---|---|---|---|
| CD40L | | # | # | HGF | | # | # | MIP-1 alpha (CCL3) | | N/A | |
| CD30 | | N/A | | MDC | | N/A | | MIP-1 beta (CCL4) | | N/A | |
| IL-22 | | # | # | IL-18 | | # | # | MIG (CXCL9) | | N/A | |
| TSLP | | N/A | | BAFF | | N/A | | MCP-2 (CCL8) | | N/A | |
| TNF beta | | # | # | G-CSF (CSF-3) | | N/A | | MMP-1 | | N/A | |
| M-CSF | | # | # | Eotaxin (CCL11) | | N/A | | IL-9 | | # | # |
| Fractalkine (CX3CL1) | | N/A | | I-TAC (CXCL11) | | N/A | | IL-2 | | # | # |
| NGF beta | | # | # | TNF-RII | | N/A | | GM-CSF | | | # |
| SCF | | # | # | MIP-3 alpha (CCL20) | | N/A | | IL-3 | | N/A | |
| IL-4 | | N/A | | MCP-1 (CCL2) | | # | # | MIP-3 alpha (CCL20) | | N/A | |
| IL-20 | | N/A | | IL-9 | | # | # | IL-27 | | N/A | |
| BLC (CXCL13) | | N/A | | IFN alpha | | | # | IL-5 | | # | # |
| APRIL | | N/A | | ENA-78 (CXCL5) | # | | # | IL-1 alpha | | # | # |
| IL-23 | | # | # | GRO alpha (CXCL1) | | # | # | SDF-1 alpha | | # | # |
| TRAIL | | # | # | VEGF-A | | # | # | IL-18 | | | # |
| IL-31 | | # | # | IL-10 | | # | | IL-10 | | # | |

**Table S3** Table of thermodynamic values used in evaluating mismatch cost in Nanoligomer algorithm. $T_M$ [2] and $K_D$[3] values were empirically determined .

| Mismatch (PNA-RNA) | $K_D$ (nM) | $T_M$ (°C) |
|---|---|---|
| None | 6.8 | 72.8 |
| A-C | 170 | 62.8 |
| A-G | 31 | 62.4 |
| A-A | 40 | 60.6 |
| C-C | 710 | 52.2 |
| C-U | 420 | 53.6 |
| C-A | 320 | 53 |
| T-U | 40 | 60 |
| T-C | 77 | 58.6 |
| T-G | 26 | 63 |
| G-G | 460 | 56 |
| G-U | 31 | 61 |
| G-A | 200 | 54.8 |

**Table S4.** Primers used for mouse tissue qPCR.

| Target | Forward 5’→3’ | Reverse 5’→3’ |
|---|---|---|
| ActB | CCACTGTCGAGTCGCGT | CGCAGCGATATCGTCATCCAT |
| L32 | CCATCTGTTTTACGGCATCATG | TGAACTTCTTGGTCCTCTTTTTGA |
| CSF2 | CAAAGAAGCCCTGAACCTCCT | GGTGAAATTGCCCCGTAGAC |
| EPO | GCCCTGCTAGCCAATTCCT | GTGGTATCTGGAGGCGACAT |

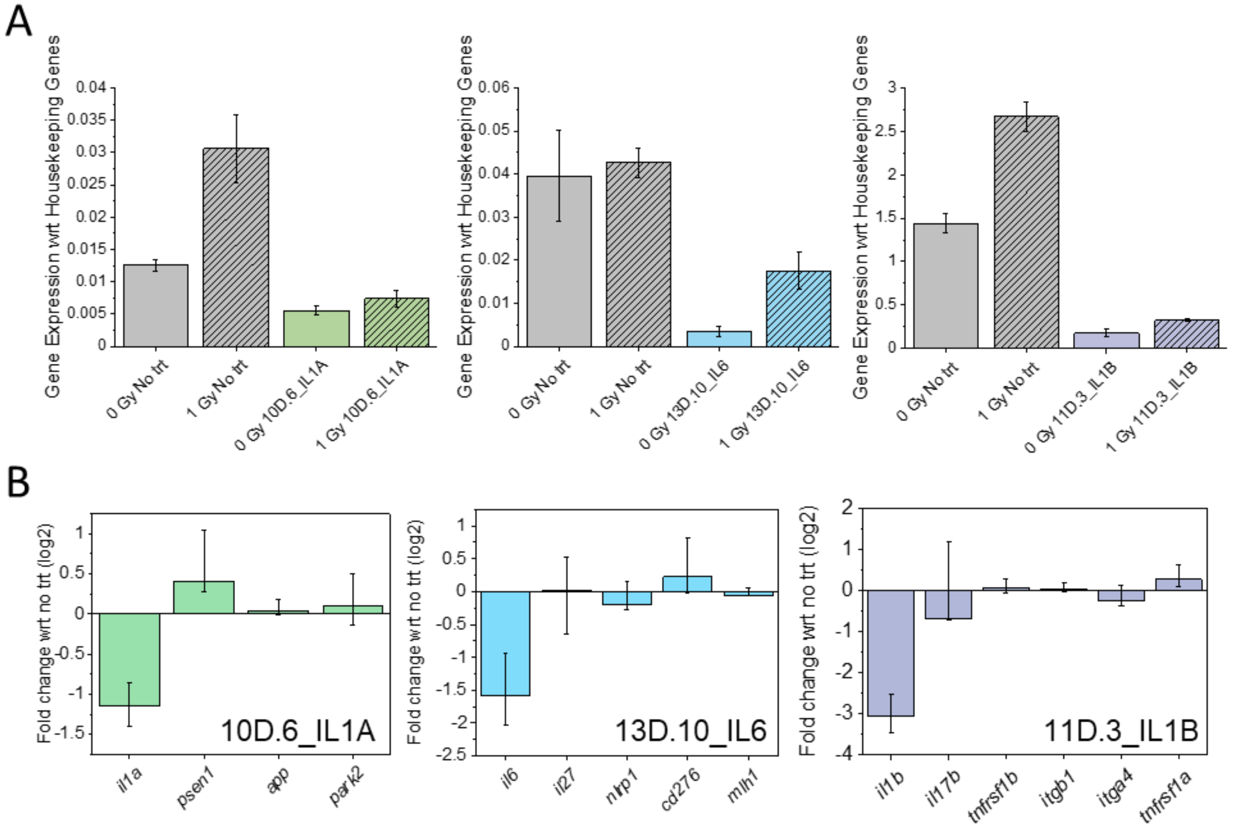


**Fig. S1 A)** Gene expression data from Quantigene analysis of PBMC after 24 h of Nanoligomer treatment and respective radiation treatment. **B)** Multiplex gene expression measurements during Nanoligomer treatment reveal no change in expression of non-pathway[4] gene as an indication of gene target specificity.[5,6]

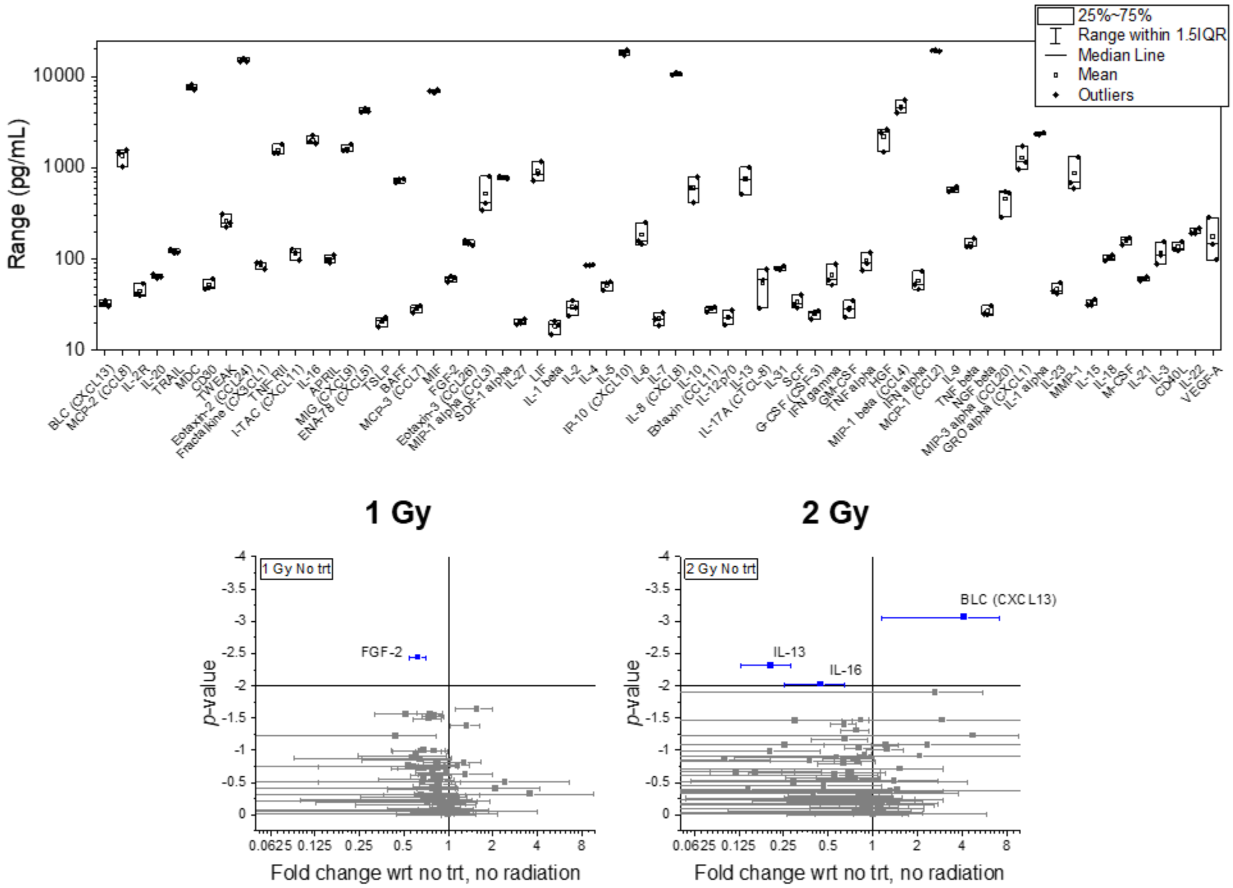


**Fig. S2** Procartaplex cytokine levels for 0 Gy no treatment PBMCs (top) and enrichment compared to no treatment of PBMC after 24 h of Nanoligomer treatment and respective radiation treatment (bottom left and right).

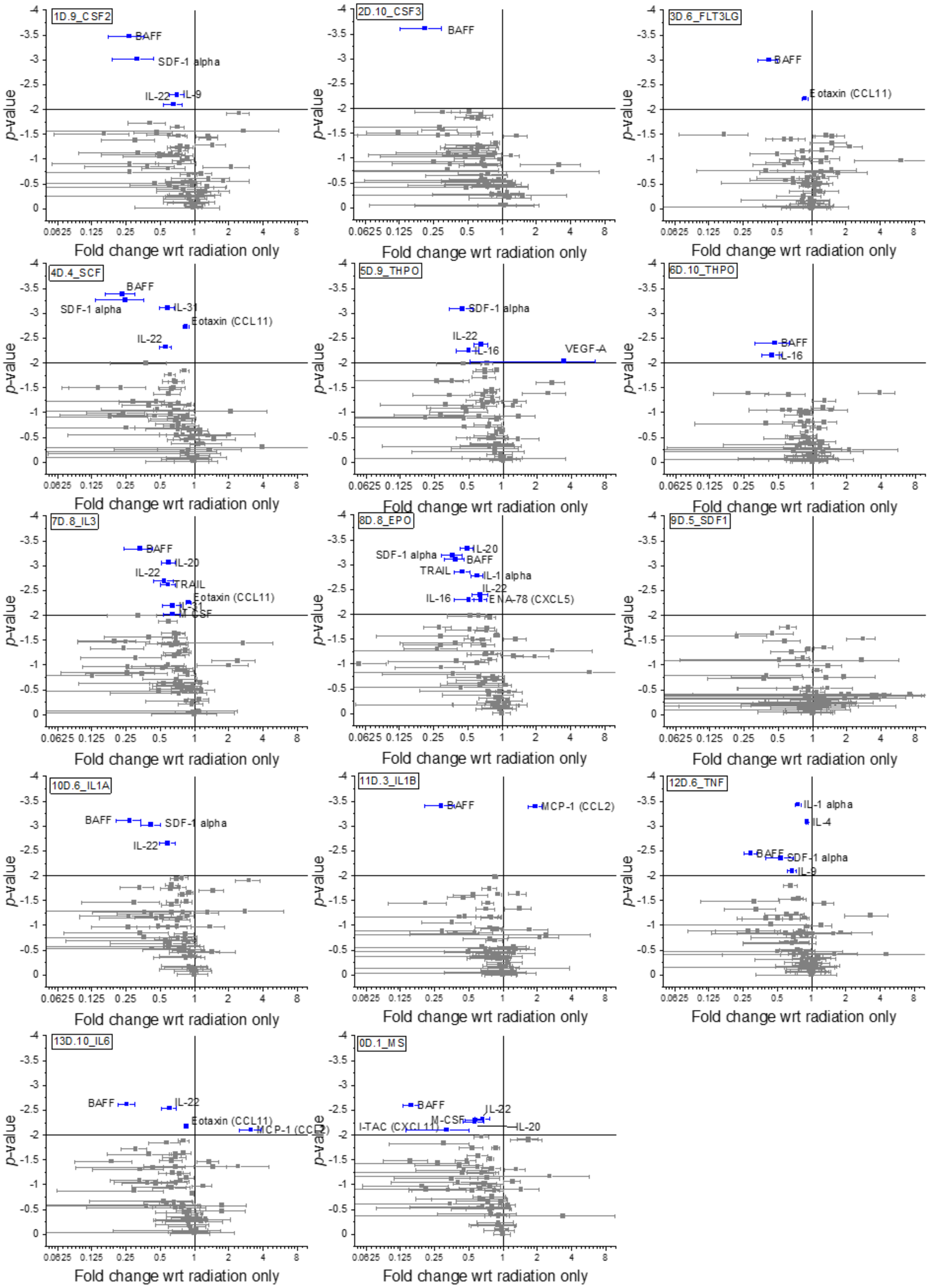


**Fig. S3** Procartaplex cytokine enrichment compared to no treatment of PBMC after 24 h of respective Nanoligomer.

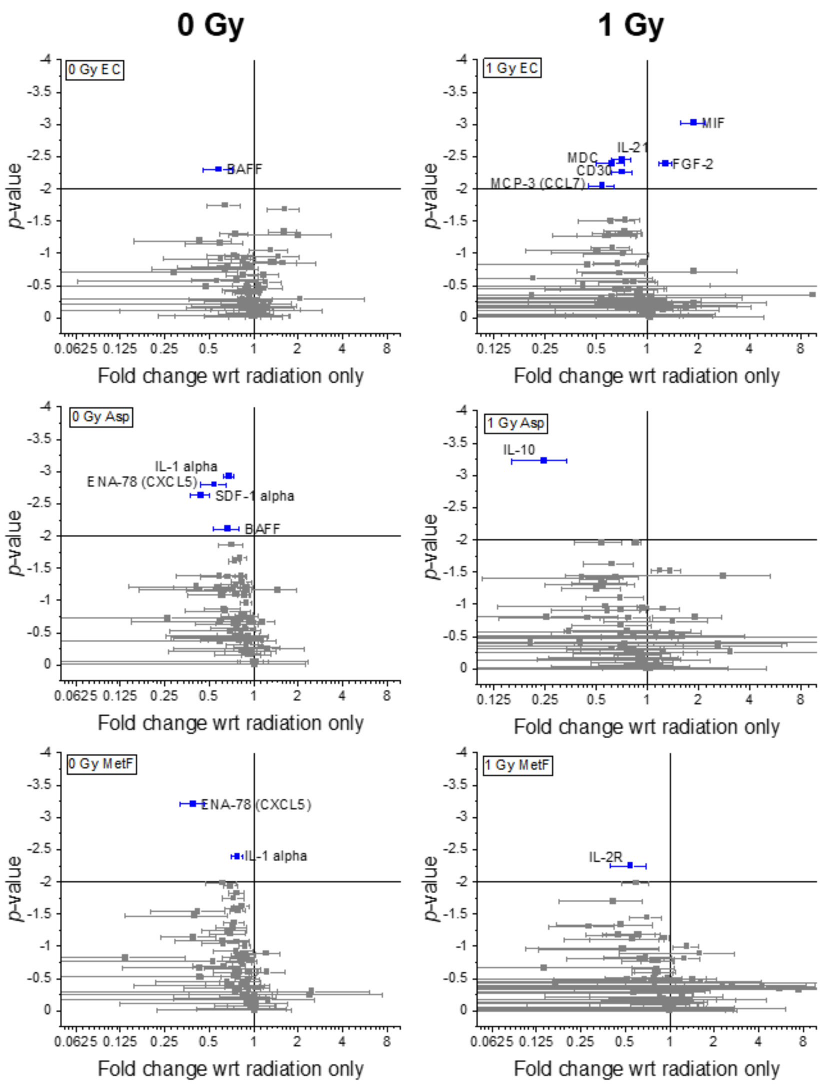


**Fig. S4** Procartaplex cytokine enrichment compared to no treatment of PBMC after 24 h of respective small molecule and respective radiation treatment.

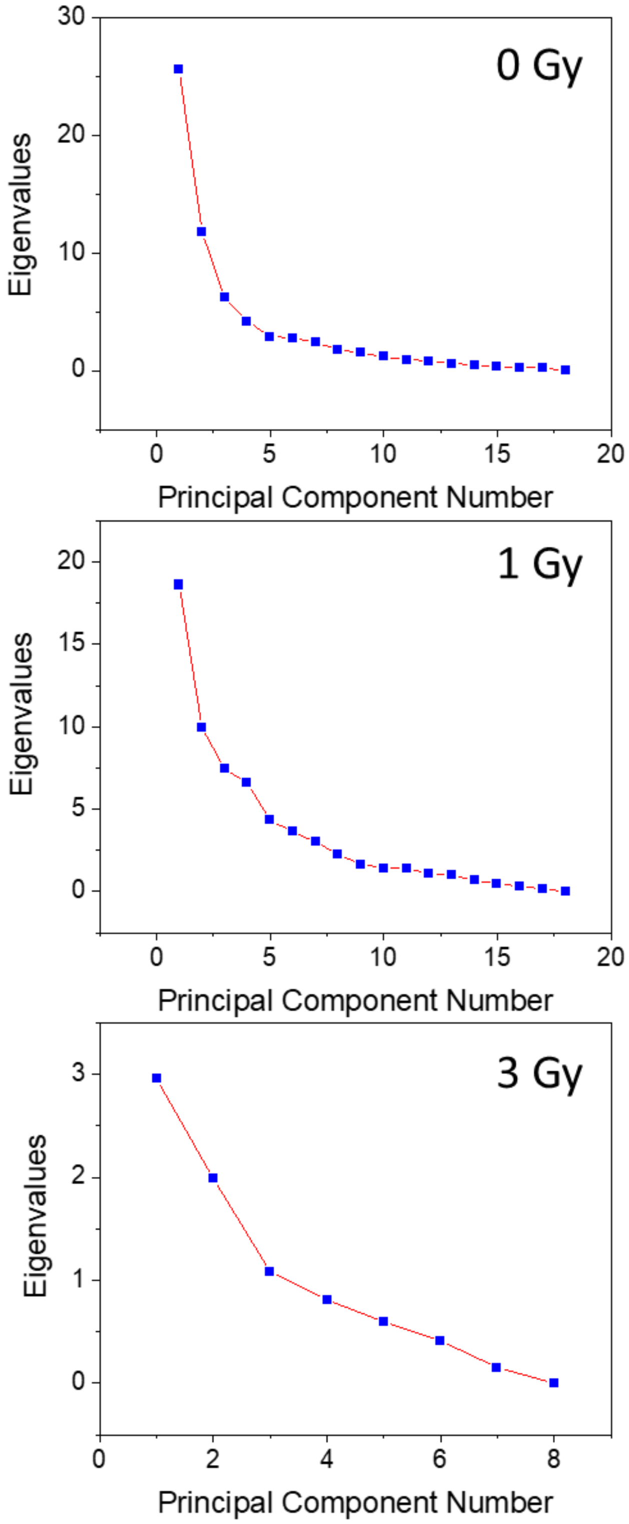


**Fig. S5** Scree plots of principal component analysis for 0 Gy cytokines panel (top), 1 Gy cytokine panel (middle), and 3 Gy cytokine panel (bottom).

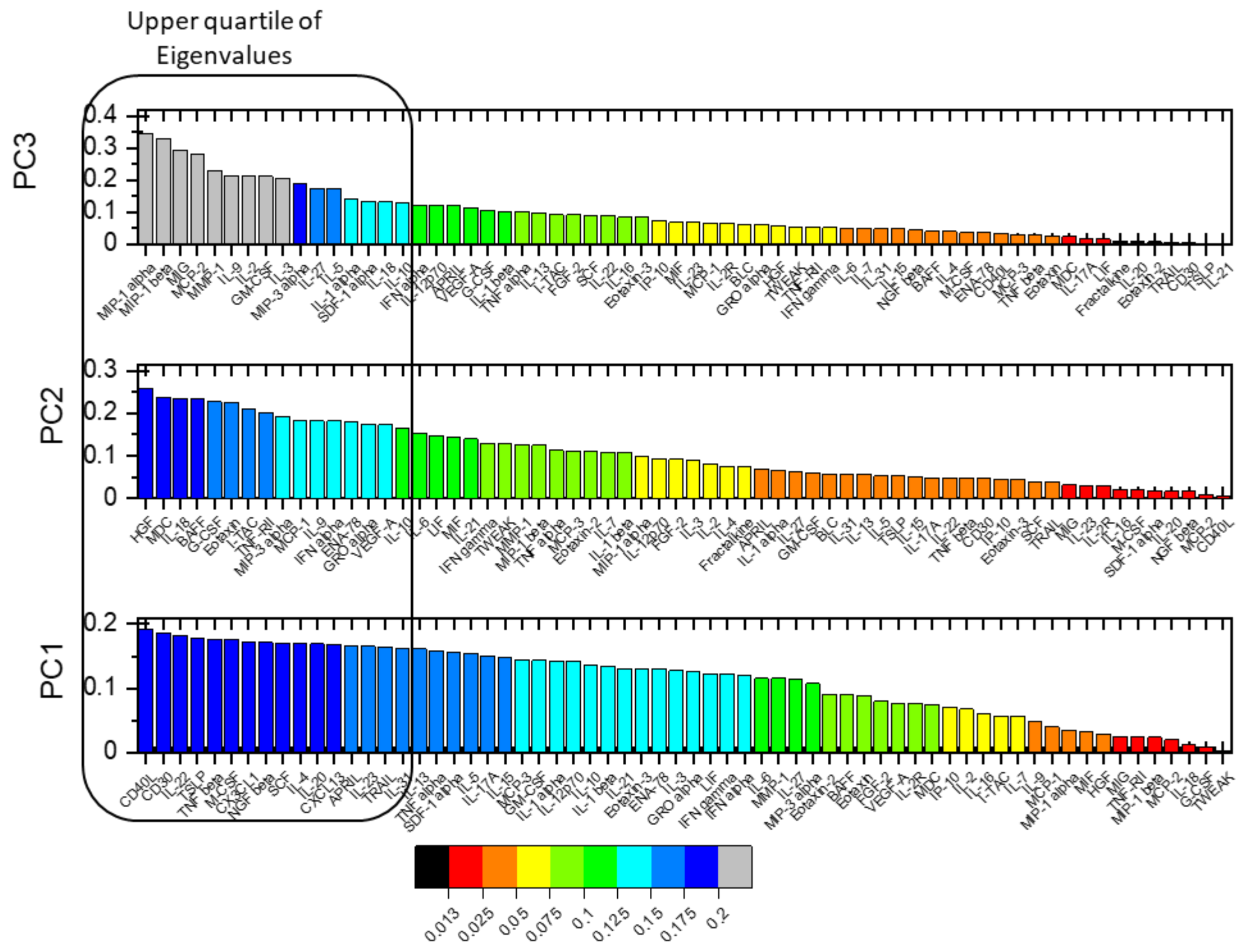


**Fig. S6** Absolute value of Eigenvalues from principal component analysis of 0 Gy data. The magnitude of the Eigenvalue corresponds to its contribution to the principal component indicated on the Y axis. The upper quartile of cytokines with the most impact on the respective principal component is shown in Table S2.

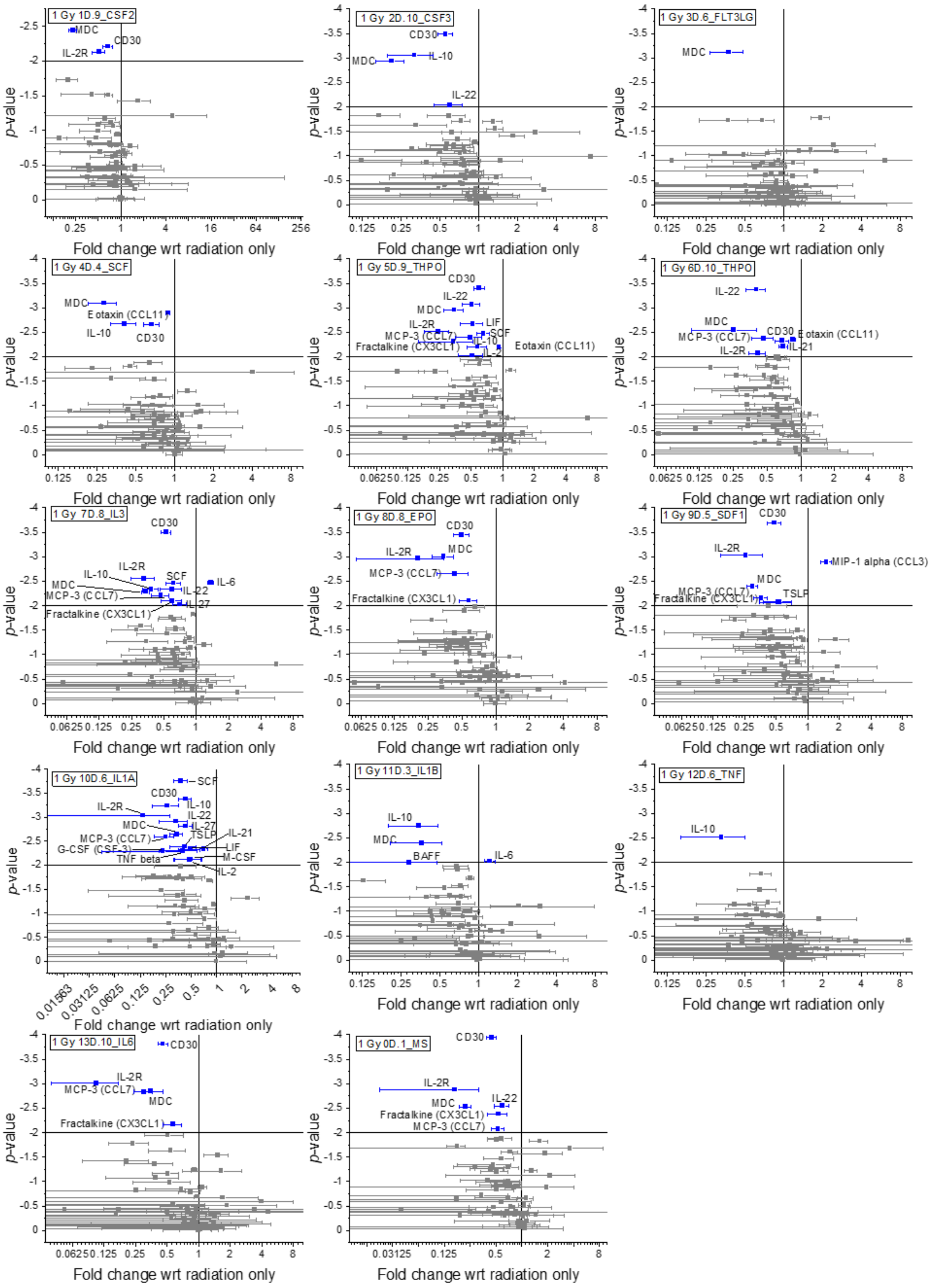


**Fig. S7** Procartaplex cytokine enrichment compared to no treatment of PBMC after 24 h of respective Nanoligomer and 1 Gy of radiation.

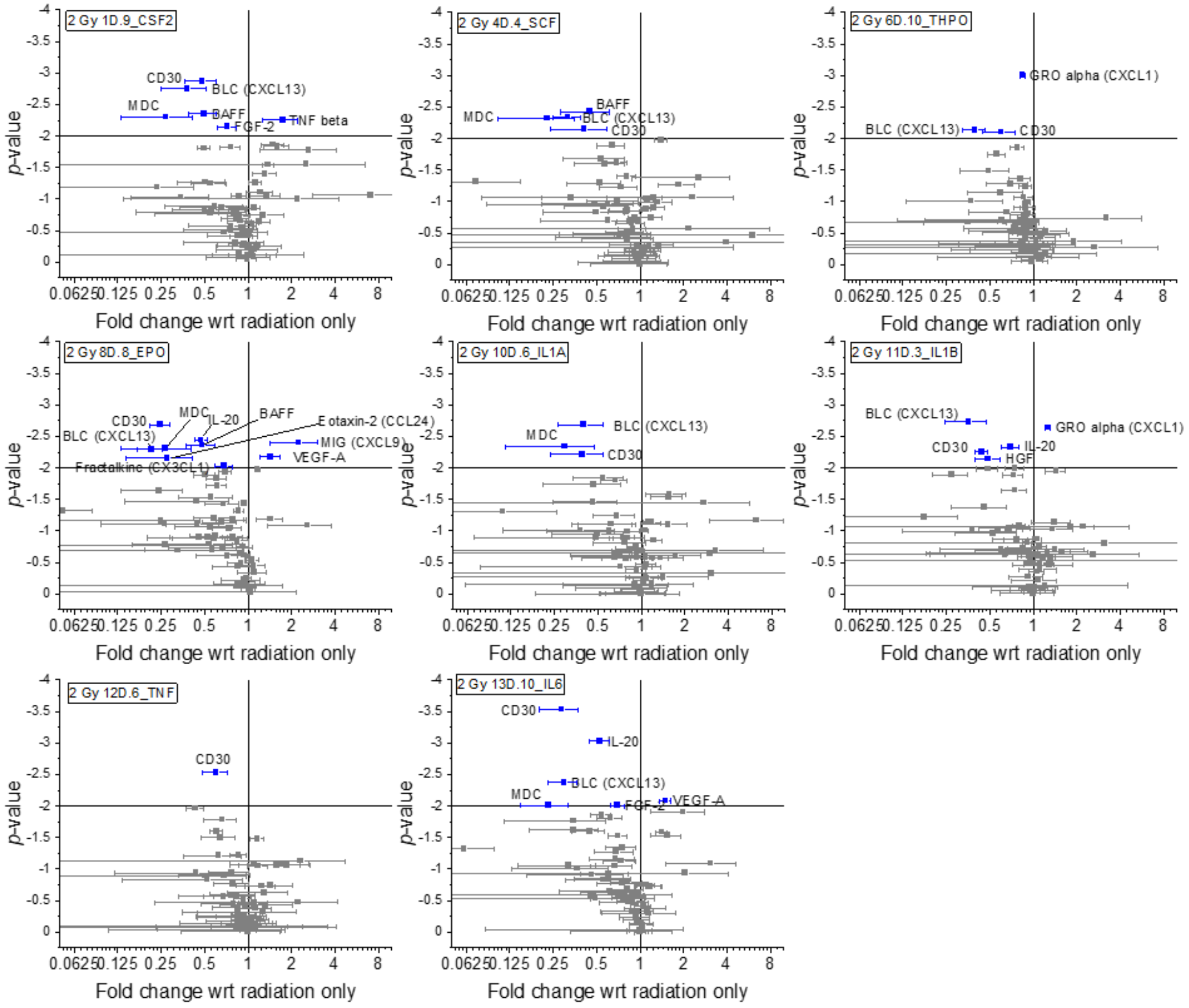


**Fig. S8** Procartaplex cytokine enrichment compared to no treatment of PBMC after 24 h of respective Nanoligomer and 2 Gy of radiation.

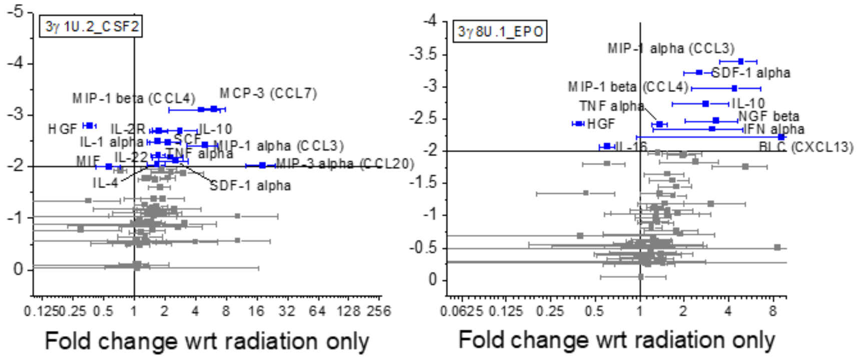


**Fig. S9** Procartaplex cytokine enrichment compared to no treatment of PBMC after 24 h of respective Nanoligomer and 3 Gy of radiation.

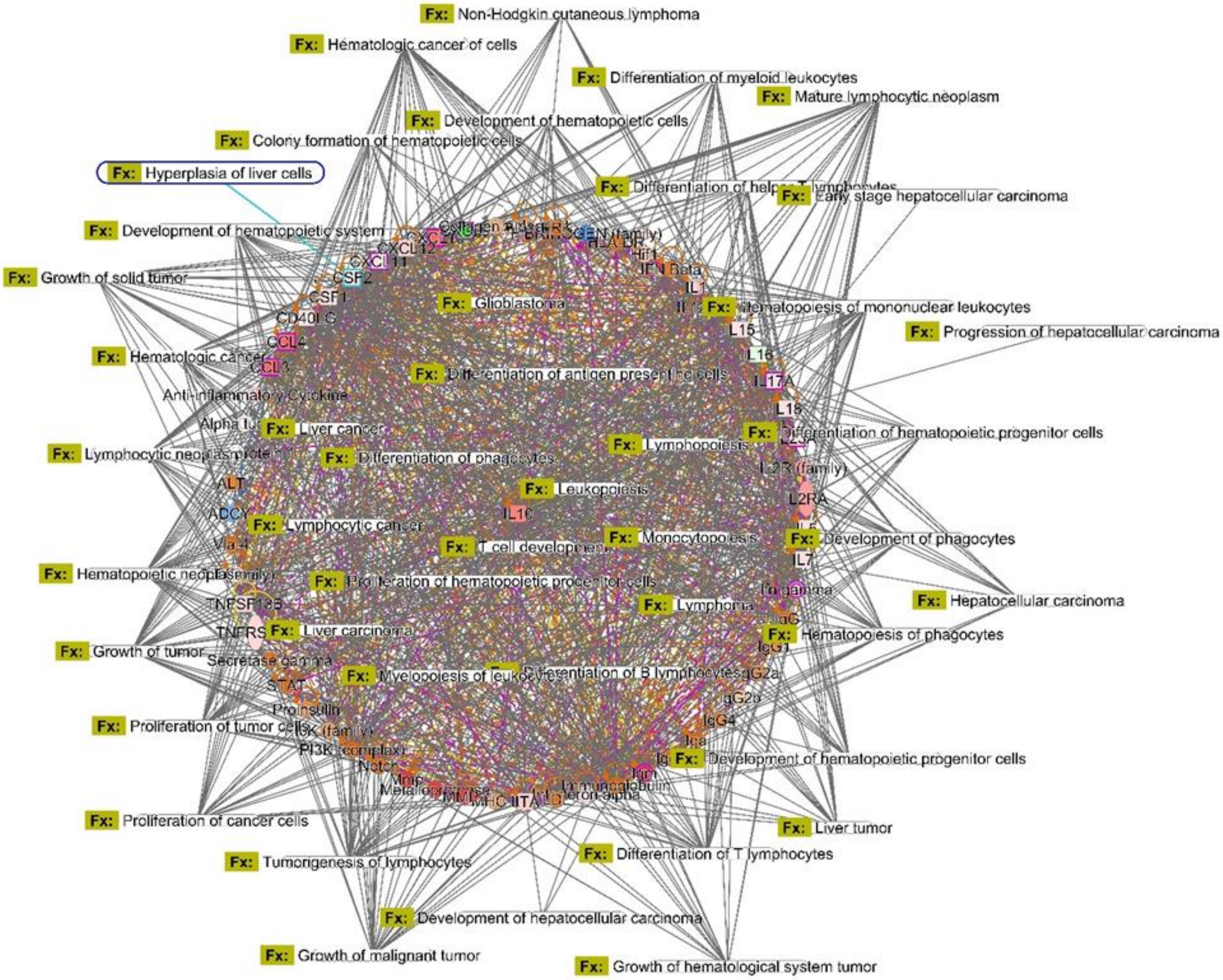


**Fig. S10.** Sachi Learn module demonstrates the differentially regulated cytokines, chemokines, growth factors, and other associated upstream regulators affect canonical pathways and other downstream expression of networks that can prevent a range of cancers, auto-immune, and other diseases that occur due to immunological dysfunction. Using the Pathway analysis shown here, Sachi Nanoligomers uniquely target disease states with divergent canonical pathways (e.g., IL-10 and pro-inflammatory cytokines). As shown here, EPO and GM-CSF activators upregulates growth factors to treat hemopoiesis while upregulating cytokines to check growth of cancer, by truly reversing the immune dysfunction induced by radiation. It is important to note that simple increase in pro-inflammatory cytokines was not the proposed objective, since such inflammatory response can induce pathology and even death, as seen recently in SARS-CoV2 (or CoVID-19 patients). However, an increase in haemopoietic proteins, proinflammatory cytokines, along with regulatory cytokines such as IL-10, an increase in Tregulatory and maintain the balance of T-cells, leads to true reversal to healthy state

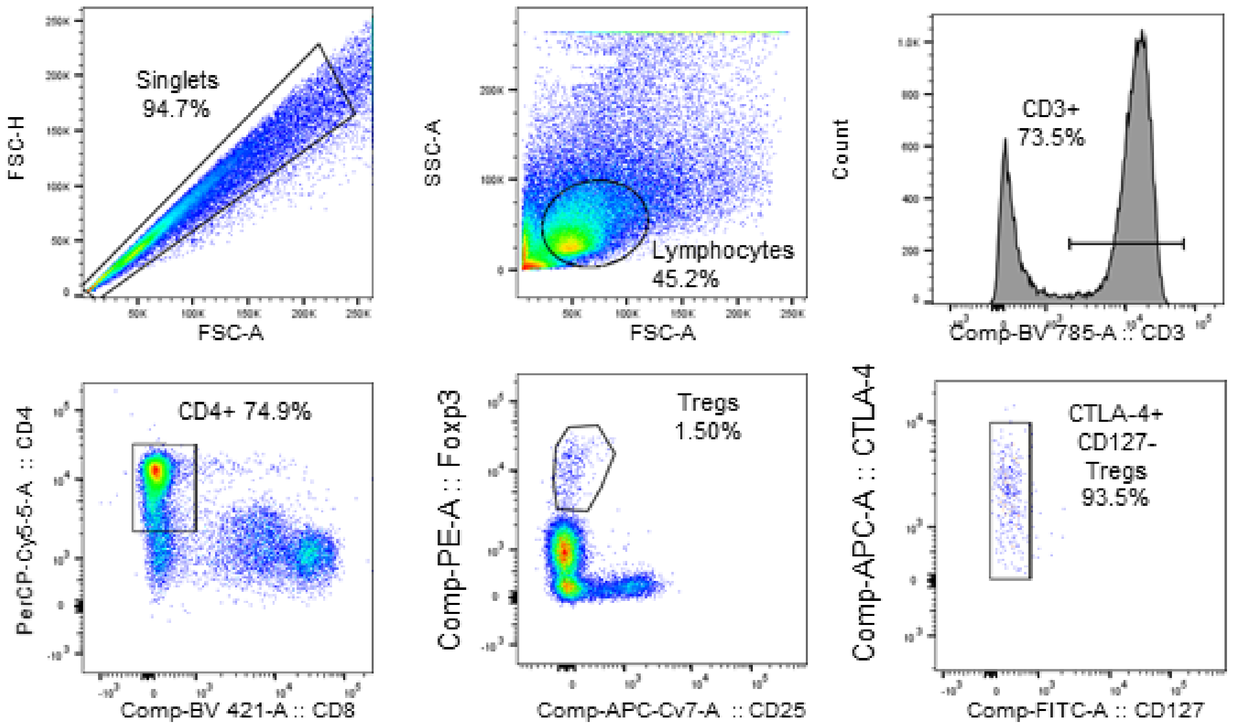


**Fig. S11** Representative flow cytometry plots showing gating scheme. Events were first gated to select singlets and then gates for lymphocytes. Antibody labeling (see Methods) was then used to select for CD3+, CD4+CD6-, followed by Foxp3+CD25+, and finally CTLA4+CD127- Tregs.

TBD Fig. S12 lung biodistribution data
*Adapted data

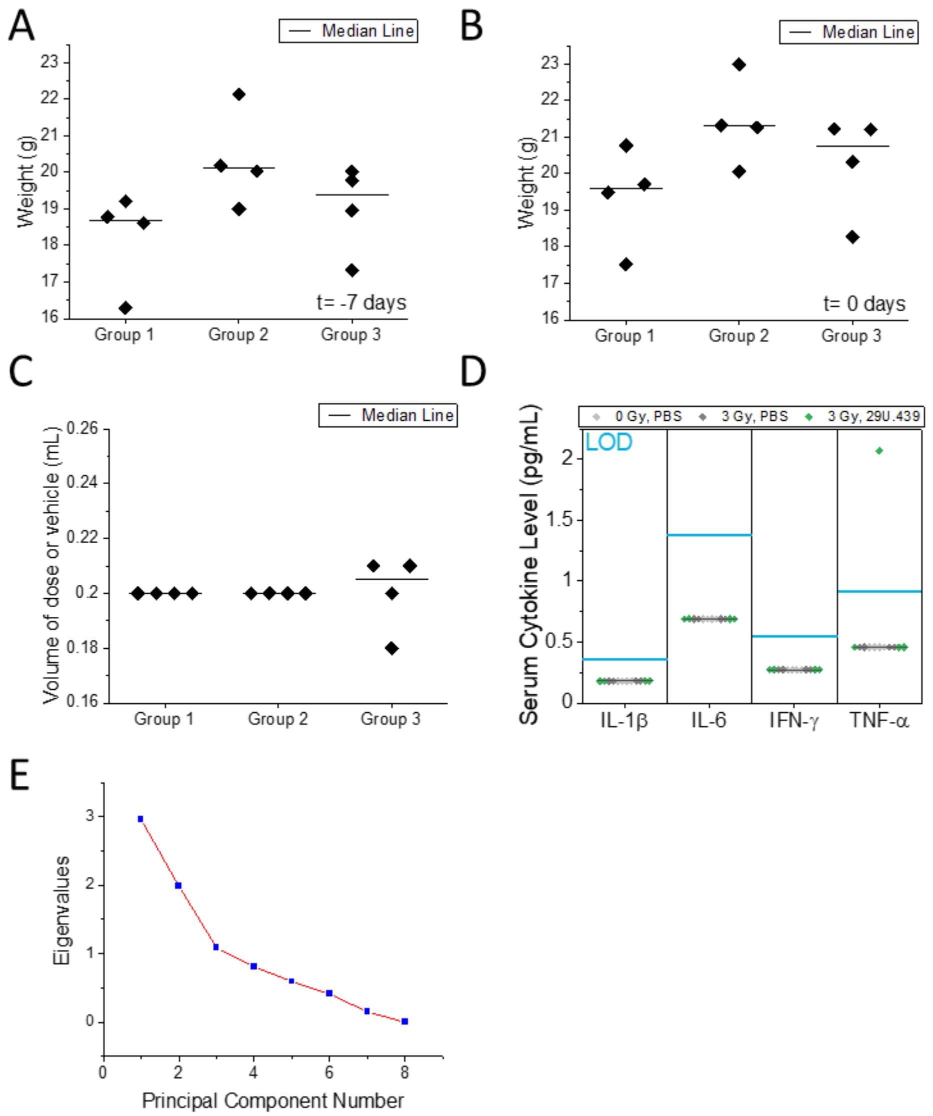


Fig. S13. A) Weights of mice 7 days before administration of Nanoligomers. B) Weight of mice on day of Nanoligomer administration. C) Volume of Nanoligomer administered via intraperitoneal injection to mice based on weight for consistent mg/kg dosage. D) Four markers were measured in murine blood serum to evaluate systemic immune response to Nanoligomers. Levels of IL-1β, IL-6, and IFN-γ were all below the limit of detection. One animal in the 3 Gy, 39U.439_EPO had TNF-α level slightly above the LOD. E) Scree plot for PCA of mouse lung cytokine expression showing importance of PC1, PC2, and PC3.

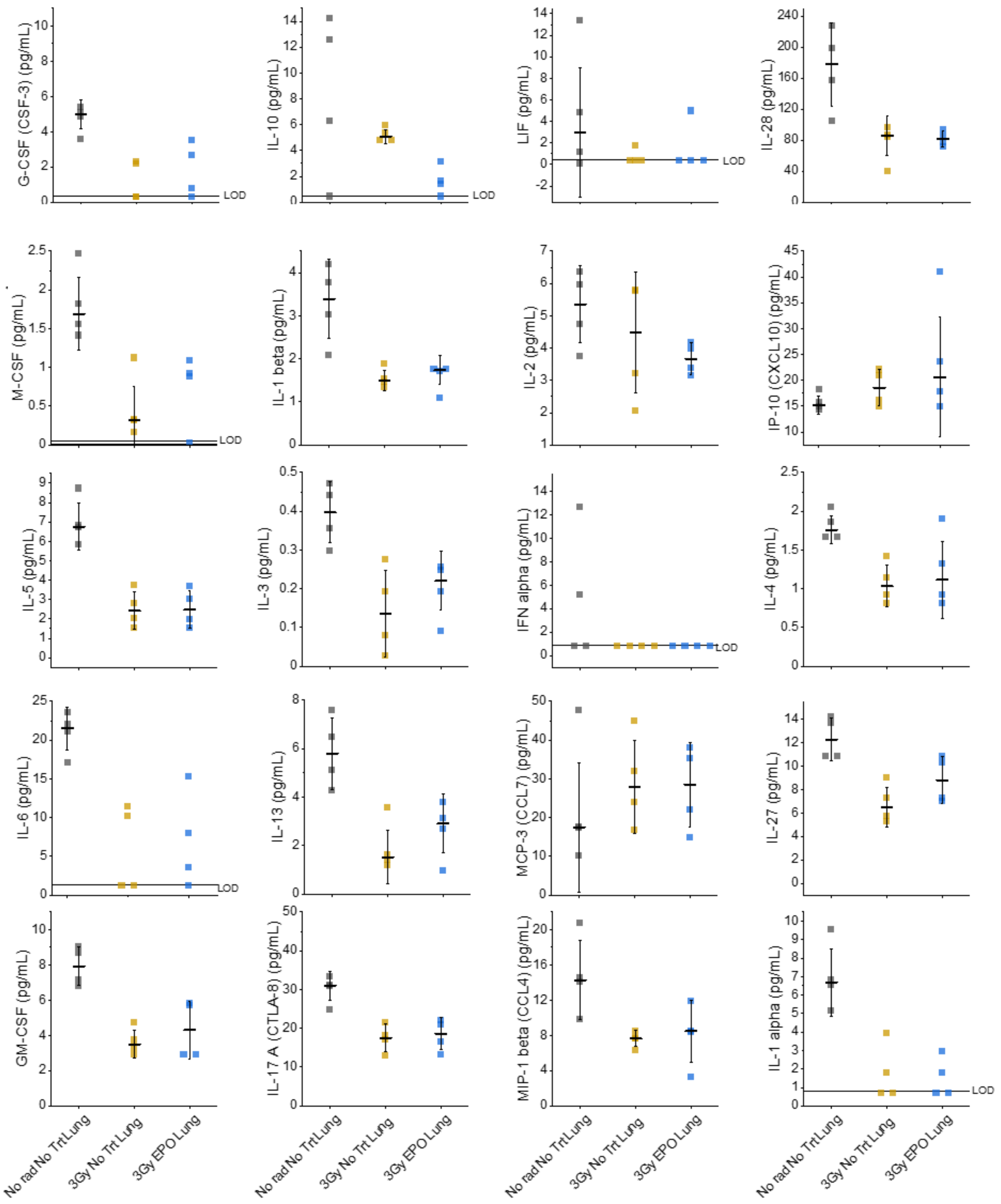


Fig. S14. Cytokine 36-plex panel results for mouse lung tissues with respective treatment.

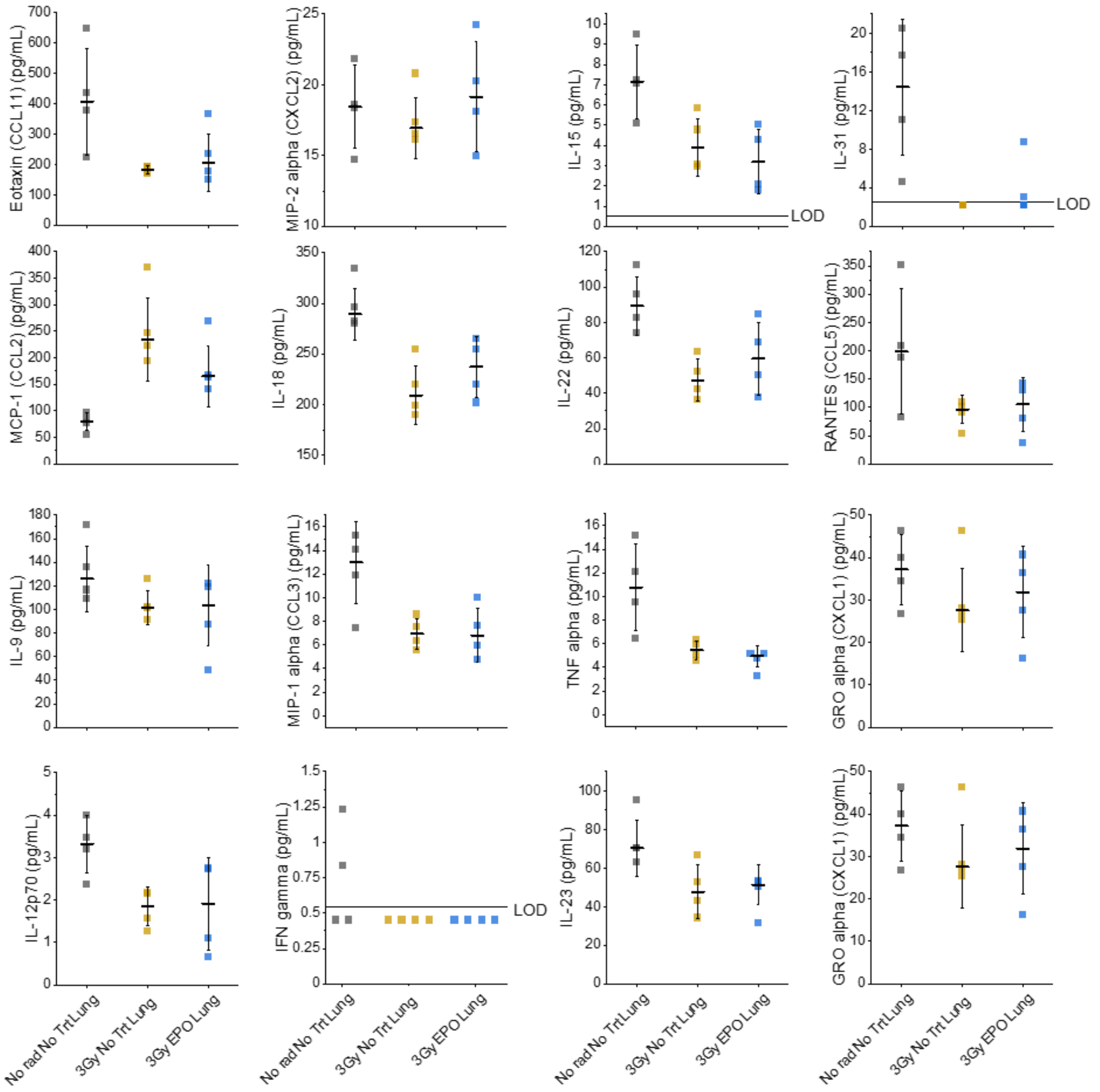


Fig. S15. Cytokine 36-plex panel results for mouse lung tissues with respective treatment.

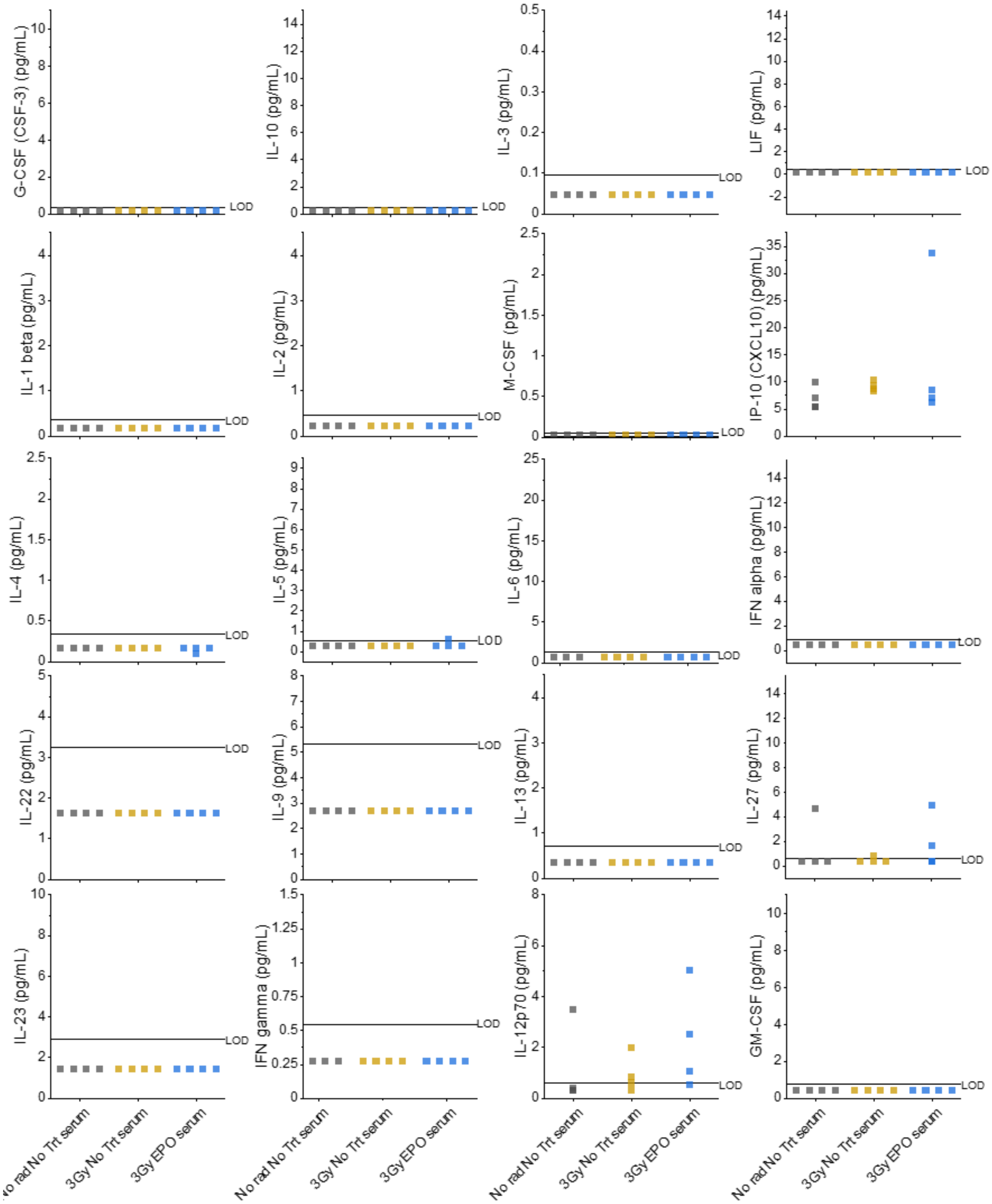

Fig. S16. Cytokine 36-plex panel results for mouse serum with respective treatment.

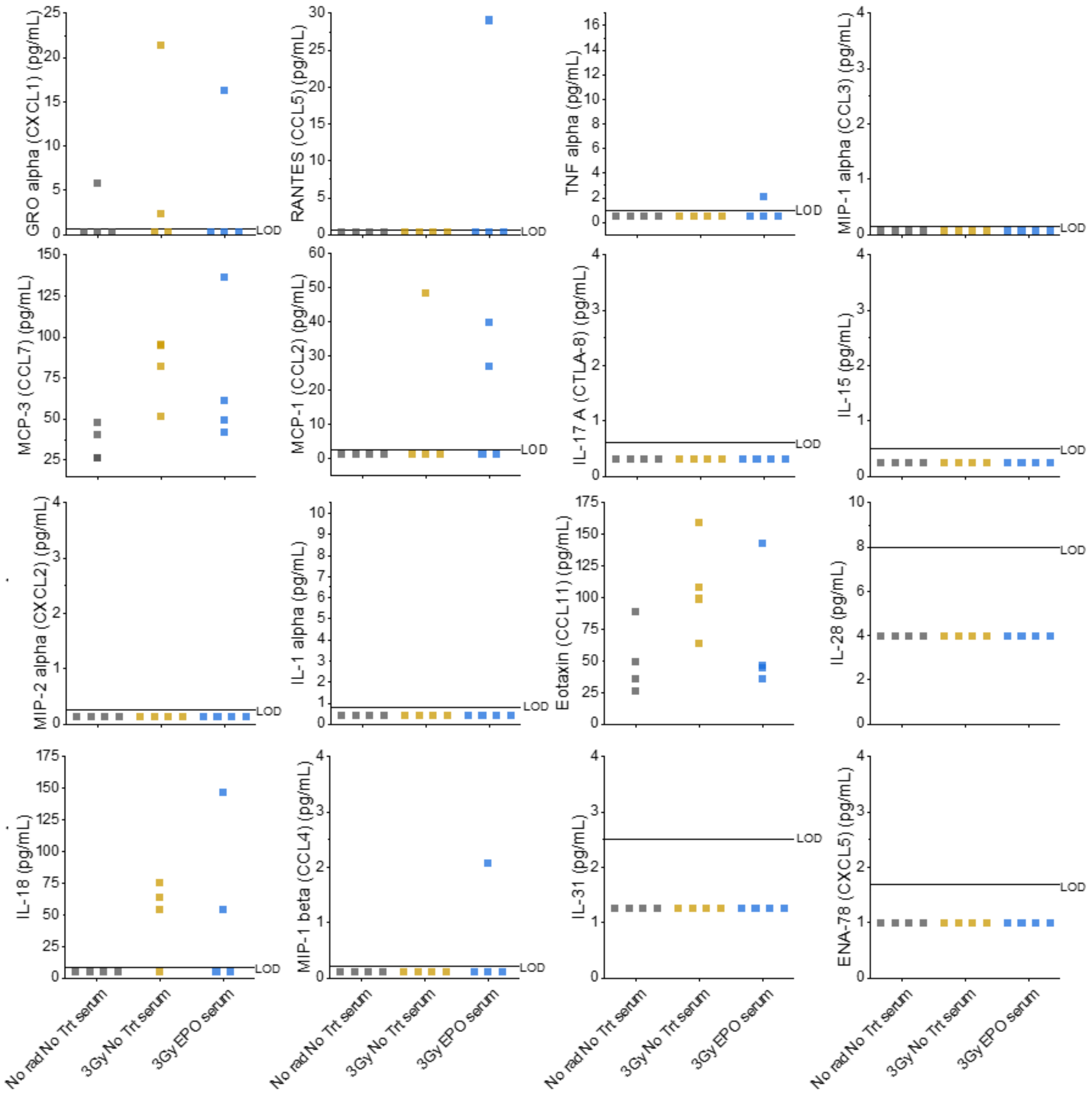


Fig. S17. Cytokine 36-plex panel results for mouse serum with respective treatment.

Supplementary References


1. Garrett-Bakelman, F. E. *et al.* The NASA twins study: A multidimensional analysis of a year-long human spaceflight. *Science (80-. ).* (2019) doi:10.1126/science.aau8650.
2. SantaLucia, J. A unified view of polymer, dumbbell, and oligonucleotide DNA nearest-neighbor thermodynamics. *Proc. Natl. Acad. Sci. U. S. A.* **95**, 1460–1465 (1998).
3. Kilså Jensen, K., Ørum, H., Nielsen, P. E. & Nordén, B. Kinetics for hybridization of peptide nucleic acids (PNA) with DNA and RNA studied with the BIAcore technique. *Biochemistry* (1997) doi:10.1021/bi9627525.
4. Szklarczyk, D. *et al.* STRING v10: Protein-protein interaction networks, integrated over the tree of life. *Nucleic Acids Res.* **43**, (2015).
5. Buehler, E. *et al.* siRNA off-target effects in genome-wide screens identify signaling pathway members. *Sci. Rep.* **2**, (2012).
6. Caffrey, D. R. *et al.* Sirna off-target effects can be reduced at concentrations that match their individual potency. *PLoS One* **6**, (2011).
